# Comparative analysis of gonadal transcriptomes between turtle and alligator identifies common molecular cues activated during the temperature-sensitive period for sex determination

**DOI:** 10.1101/2023.06.09.544319

**Authors:** Kenji Toyota, Hiroshi Akashi, Momoka Ishikawa, Katsushi Yamaguchi, Shuji Shigenobu, Tomomi Sato, Anke Lange, Charles R. Tyler, Taisen Iguchi, Shinichi Miyagawa

## Abstract

The mode of sex determination in vertebrates can be categorized as genotypic or environmental. In the case of genotypic sex determination (GSD), the sexual fate of an organism is determined by the chromosome composition with some having dominant genes, named sex-determining genes, that drive the sex phenotypes. By contrast, many reptiles exhibit environmental sex determination (ESD), whereby environmental stimuli drive sex determination, and most notably temperature. To date, temperature-dependent sex determination (TSD) has been found in most turtles, some lizards, and all crocodilians, but commonalities in the controlling processes are not well established. Recent innovative sequencing technology has enabled investigations into gonadal transcriptomic profiles during temperature-sensitive periods (TSP) in various TSD species which can help elucidate the controlling mechanisms. In this study, we conducted a time-course analysis of the gonadal transcriptome during the male-producing temperature (26L) of the Reeve’s turtle (Chinese three-keeled pond turtle) *Mauremys reevesii*. We then compared the transcriptome profiles for this turtle species during the TSP with that for the American alligator *Alligator mississippiensis* to identify conserved reptilian TSD-related genes. Our transcriptome-based findings provide an opportunity to retrieve the candidate molecular cues that are activated during TSP and compare these target responses between TSD and GSD turtle species, and between TSD species.

**Highlights:**

- A time-course gonadal RNA-seq was conducted using *Mauremys reevesii*.
- Sexual differentiation genes in males were activated at an earlier stage than the ones in females.
- Turtle-alligator comparative analysis revealed novel candidate TSD genes.

## 1. Introduction

Sex determination in vertebrates can be broadly categorized into genotypic or environmental. In the case of genotypic sex determination (GSD), the sexual fate of an organism is determined by the sex chromosomes inherited from both parents, with some having dominant genes on heterozygous sex chromosomes (such as *sex-determining region Y* (*Sry*) in mice) (Koopman et al., 1991), and others having dose-sensitive genes on homozygous sex chromosomes (such as *doublesex and mab–3 related transcription factor 1* (*DMRT1*) in the chick) (Smith et al., 2009). However, many reptiles and some fish exhibit environmental sex determination (ESD), where incubation temperature during embryonic development in particular is a key environmental cue for their sex determination system (Capel 2017; Nagahama et al., 2021). To date, temperature-dependent sex determination (TSD) has been shown in reptile species including most turtles, some lizards, and all crocodilians studied, but not snakes (Augstenová et al., 2018; Nagahama et al., 2021; Rovatsos et al., 2015). Although still controversial, GSD and TSD systems are not necessarily exclusive, but rather these systems may be part of a continuum because several reptiles, such as the central bearded dragon *Pogona vitticeps*, have been shown to exhibit intermediate states, illustrating that the sex-determining signaling cascade governed by sex-linked factors can be overridden by thermal-induced sex reversal (Quinn et al., 2007; Whiteley et al., 2021).

The key event for the sexual fate decision in TSD species is the detection of environmental temperature during a specific embryonic period known as the temperature-sensitive period (TSP). TSP has been investigated in many TSD species by egg-incubation experiments whereby egg incubation temperatures are manipulated during various embryonic developmental periods with precise developmental staging and the resulting sex of the individuals analyzed afterwards (Bull 1987; Lang and Andrews, 1994). Although the gonadal developmental stages of turtles and alligators during the TSP are similar, corresponding with the early stages at which the genital ridges develop (Wibbels et al., 1991), the temperature ranges that induce males and females are different. For example, the red-eared slider turtle *Trachemys scripta* becomes a male when incubated at low temperatures (26°C) and a female when incubated at high temperatures (31°C) (Crews et al., 1994). While, in the case of the American alligator *Alligator mississippiensis* females arise at a temperature of 30°C (or lower) and males at a higher temperature (33°C) (Ferguson and Joanen, 1982). After the TSP when the sexually dimorphic gonads have developed, the sex is no longer reversible regardless of the changes in incubating temperature.

Recent innovative sequencing technology has enabled the investigation of the gonadal transcriptomic profiles during TSP in various TSD species, such as *T*. *scripta* (Czerwinski et al., 2016), the painted turtle *Chrysemys picta* (Radhakrishnan et al., 2017), and *A*. *mississippiensis* (Yatsu et al., 2016). In this study, we carried out a time-course analysis of the gonadal transcriptome during the temperature-sensitive period (TSP) in the Chinese three-keeled pond turtle *Mauremys reevesii* at the male-producing temperature (MPT) of 26°C. *M. reevesii* is a freshwater turtle distributed across China, the Korean Peninsula, and Japan (Rhodin et al., 2017; Yin et al., 2016). It is an endangered species in China, but is commercially produced for aquaculture in China and Japan. To date, *T*. *scripta*, has been the turtle model of choice for studies into TSD in turtles, however, it has become an invasive alien species in Asia and because of this is becoming less favored as a study model (Lowe et al., 2020). As a consequence, *M. reevesii* is attracting attention as a new model species for TSD research particularly in the Asian region. Recently, its genome has been sequenced with a chromosome-level assembly (Liu et al., 2021) and a precise embryonic developmental staging, including a description of its gonadal development have been established (Akashi et al., 2022), paving the way for understanding the gene cascades underlying embryogenesis and gonadal developmental processes during TSP in this species. Here we compare the transcriptome profiles during TSP between *M*. *reevesii* and *A. mississippiensis* (Yatsu et al., 2016) to identify the conserved reptilian TSD-related genes, with the prospect of identifying the candidate molecular clues activated during TSP and for comparing the differences in the molecular events controlling sex determination between TSD and GSD in turtle species, and between TSD species more widely in reptiles.

## 2. Materials and methods

### 2.1 Sample collection and experimental design

Turtle eggs of the Chinese three-keeled pond turtle *Mauremys reevesii* were purchased from Kondo farm in Maniwa, Okayama, Japan between June and August in 2016 and 2017. Eggs were collected from nests 1–3 days after oviposition, transported to the Tokyo University of Science laboratory, and incubated at 26[(MPT for this species). The rate of embryonic development was predicted based on previous data and staged according to those criteria (see: Akashi et al., 2022; Greenbaum 2002). All eggs were incubated under MPT conditions until embryonic developmental stage 15, at which point eggs were randomly split into two incubating temperatures, MPT and female-producing temperature (FPT; 31[) (Figure 1A). This design (incubation at a lower temperature until TSP) was adopted in the previous RNA sequencing (RNA-seq) analysis of the gonadal differentiation of American alligators (Yatsu et al., 2016). The gonads were sampled at embryonic stages 15, 16, 17, and 18 under MPT conditions and stage 18 under FPT. For all samplings, three biological replicates (n = 3) were prepared except for stage 18 under MPT (n = 2). Additionally, due to the poor quality of the nucleic acids for one sample, there were also only two samples (n=2) for analysis for stage 16 under MPT.

**Figure 1.**
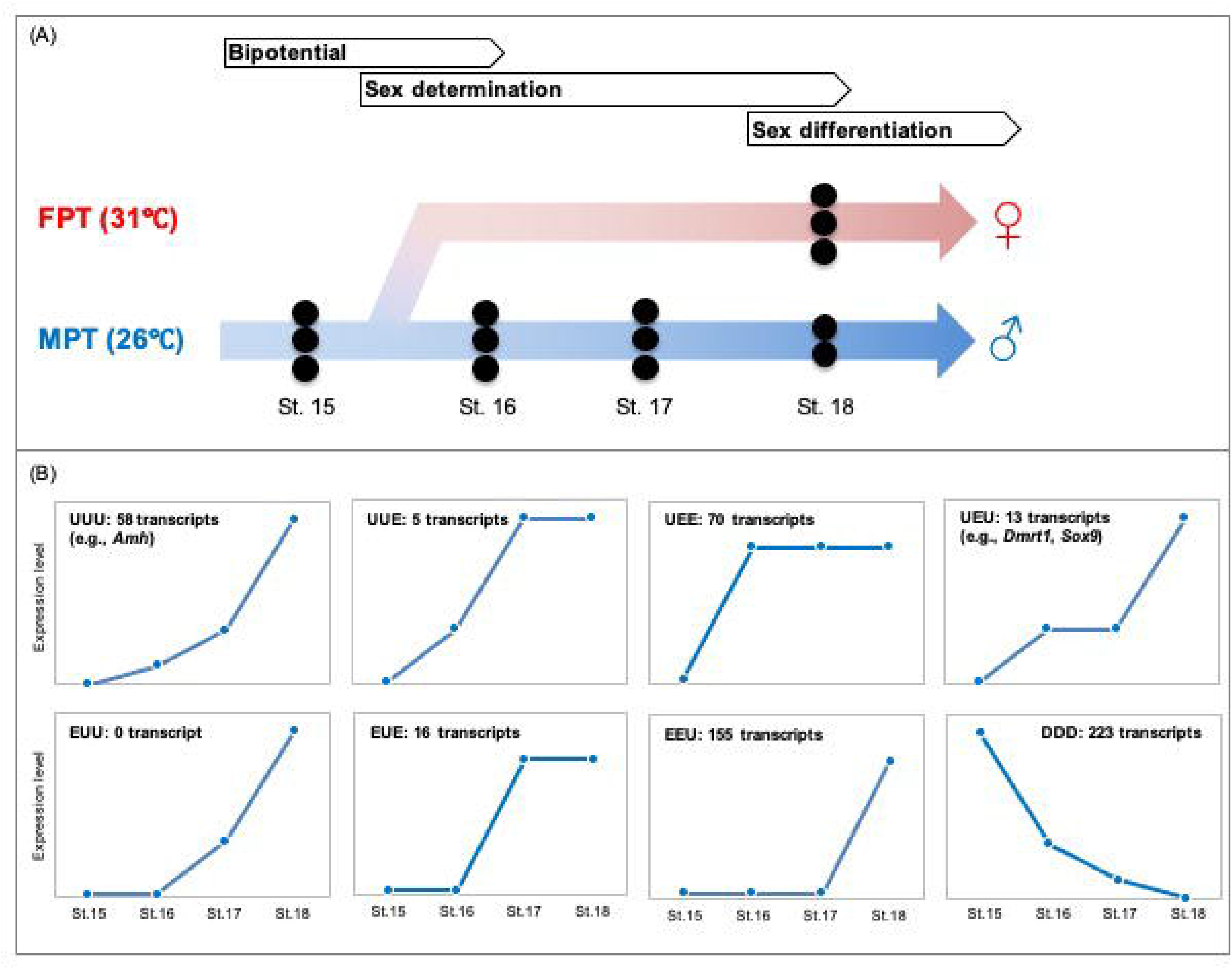
The experimental design of the RNA-seq analysis for *M*. *reevesii* is illustrated (A). Bipotential, sex determination and sex differentiation periods are indicated. Eggs were first incubated under male-producing temperature (MPT; indicated in blue) until just prior to the onset of sexual differentiation (stage 15), when they were either shifted to female-producing temperature (FPT; indicated in red) or kept at MPT. Gonadal regions were sampled from individuals at several subsequent developmental stages (stages 16, 17, and 18 under MPT and stage 18 under FPT). Models of expression patterns during developmental time-course (stages 15-18) in MPT incubated eggs (B). U, E, and D indicate the "up", "equal (no change)", and "down", respectively. For example, UUU means expression levels gradually increased depending on developmental stage progression. The annotation list of UUU, EEU, and DDD is in Table 2.

All experiments were performed in accordance with relevant animal practice guidelines and regulations that were approved by the Institutional Animal Care and Use Committee at the Tokyo University of Science. All efforts were made to minimize the suffering of the animals.

### 2.3 RNA preparation and sequencing

Total RNA from the gonad of each individual was extracted using ISOGEN reagent (Nippon Gene, Toyama, Japan) and was purified with the Promega SV Total RNA Isolation system (Promega, Madison, WI, USA) according to the manufacturer’s instructions. In addition to gonads, the following tissues were prepared for constructing a more comprehensive gene catalog: brain, liver, and urogenital from stage 14 under MPT conditions, and liver and tail from stage 15 under MPT conditions. The RNA treated with RNase-free DNase was cleaned up using the RNeasy Mini kit (Qiagen, Valencia, CA, USA) according to the manufacturer’s protocol. The quality and concentration of total RNA were validated by NanoDrop (Thermo Fisher Scientific, Waltham, MA, USA), Qubit (Life Technologies, Waltham, MA, USA), and 2100 Bioanalyzer RNA poco kit assay (Agilent Technologies, Santa Clara, CA, USA). RIN scores of all samples outputted by 2100 Bioanalyzer were more than 8.4. The libraries for transcriptome analyses were prepared from 100 ng of total RNA using the TruSeq RNA Sample Preparation v2 kit (Illumina, San Diego, CA, USA) following the manufacturer’s protocols with minor modifications (Yatsu et al., 2016). The libraries were then validated using a KAPA library quantification kit (Kapa Biosystems, Woburn, MA, USA) and 2100 Bioanalyzer High Sensitivity DNA assay (Agilent Technologies). Finally, the libraries were multiplexed into one pool and sequenced using the Illumina HiSeq2500 instrument (Illumina) at National Institute for Basic Biology in Okazaki, Japan. Sequencing was performed as 101 bp, paired-end reads in three lanes. The RNA-seq reads are available under the accession number: DRA016380.

### 2.4 Sequencing data pre-processing and *de novo* assemblies with transcript annotation

The quality of output sequences was inspected using the Fast QC program (version 0.11.2, available online at: http://www.bioinformatics.babraham.ac.uk/projects/fastqc). FastQC: a quality control tool for high throughput sequence data. Available online at: http://www.bioinformatics.babraham.ac.uk/projects/fastqc]. Illumina adapters were removed from raw reads with Cutadapt (version 1.16). Cleaned reads from the same treatment group (biological replicates) were assembled together using the RNA-seq *de novo* assembler Trinity (version 2.8.4) with the paired-end mode (Grabherr et al., 2011). Then all Trinity-assembled transcriptome (gonads: stages 15, 16, 17, 18 at MPT, stage 18 at FPT; other tissues mix composing of urogenitals, livers, brains, and tails) were merged into a single fasta file and then submitted to the EvidentialGene tr2aacds pipeline (available online at: https://sourceforge.net/projects/evidentialgene/) to generate a single assembly with minimal redundancy while maximizing the maintenance of long coding sequence regions in each contig. Finally, the tr2aacds pipeline produced the “okay” set of transcripts that were regarded as optimal. The reads from each biological replicate were mapped to the assembled transcripts for quantification by Salmon (version 0.9.1) with the “--dumpEq” option (Patro et al., 2017). Then, contigs were clustered based on the proportion of shared reads and expressions by Corset (version 1.09) (Davidson and Oshlack, 2014). Corset generated the clusters and outputs as a table of counts containing the number of reads uniquely assigned to each cluster. BLAST X was used to find matches to highly similar reference sequences in the *Mus musculus* (GRCm38.p6, Ensemble99) with AC-DIAMOND package (version 0.9.22) (Mai et al., 2018). The completeness of orthologs of the Corset-generated transcriptome was examined using BUSCO version 5.0.0 against eukaryote_odb10 (Creation date: 2020-09-10, number of species: 70, number of BUSCOs: 255). Using RNA-seq data from *A. mississippiensis* stages 22-23 (accession number: DRA004128-41, Yatsu et al., 2016), which is considered to be a temperature-sensitive period for sex determination similar to stage 18 of turtle *M*. *reevesi*, the analysis was performed by the same analytical pipeline described above to allow for comparison between both species.

### 2.5 Time-course gene expression analysis and enrichment analyses

Differential expression analysis over the developmental time course across stages 15 to 18 of *M*. *reevesii* at MPT was determined using the EBSeqHMM package (version 3.10) (Leng et al., 2015). Genes were grouped into expression paths (e.g., “UP-UP-UP”, “DOWN-DOWN-DOWN”) and were considered as differentially expressed genes if FDR < 0.05. The DAVID database (Sherman et al., 2022, last accessed: 27th April 2023) was used to conduct further gene ontology (GO) and Kyoto Encyclopedia of Genes and Genomes (KEGG) enrichment analyses using only expression profiles with at least a 25% posterior probability (Max PP).

### 2.6 Differential expression analysis

Using the Corset-generated count data, differentially expressed genes (DEGs) were calculated using the DESeq2 package in the SARTools package (version 1.6.6) (Varet et al., 2016) between MPT and FPT in *M*. *reevesi* (stage 18) and *A*. *mississippiensis* (stages 22-23). Principal component analysis was also conducted using the DESeq2 package in the SARTools package. To compare the gene expression profiles between both species, the Ensemble mouse protein ID (ENSMUSP) was assigned to both transcripts, and those with the same ID were defined as the homologous transcript. Then, a two-dimensional plot was created using logFC (fold change) values for both.

## 3. Results and discussion

### 3.1 Sequencing and transcriptome assembly

Illumina HiSeq2500 sequencing yielded a total of 336,606,198 paired-end clean reads (2 × 101 bp). The transcriptome assembly processes produced 113,460 putative transcripts using the Trinity, EvidentialGene, and Corset pipelines. Among them, 22,464 transcripts could be assigned to the publicly available *M*. *musculus* database (GRCm38.p6, Ensemble99). The final transcripts were evaluated by BUSCO for completeness based on expectations of gene content against the eukaryotic gene dataset, indicating that our *de novo* assembly pipeline built a reliable comprehensive gene catalog for *M. reevesii* (Table 1).

**Table 1.**
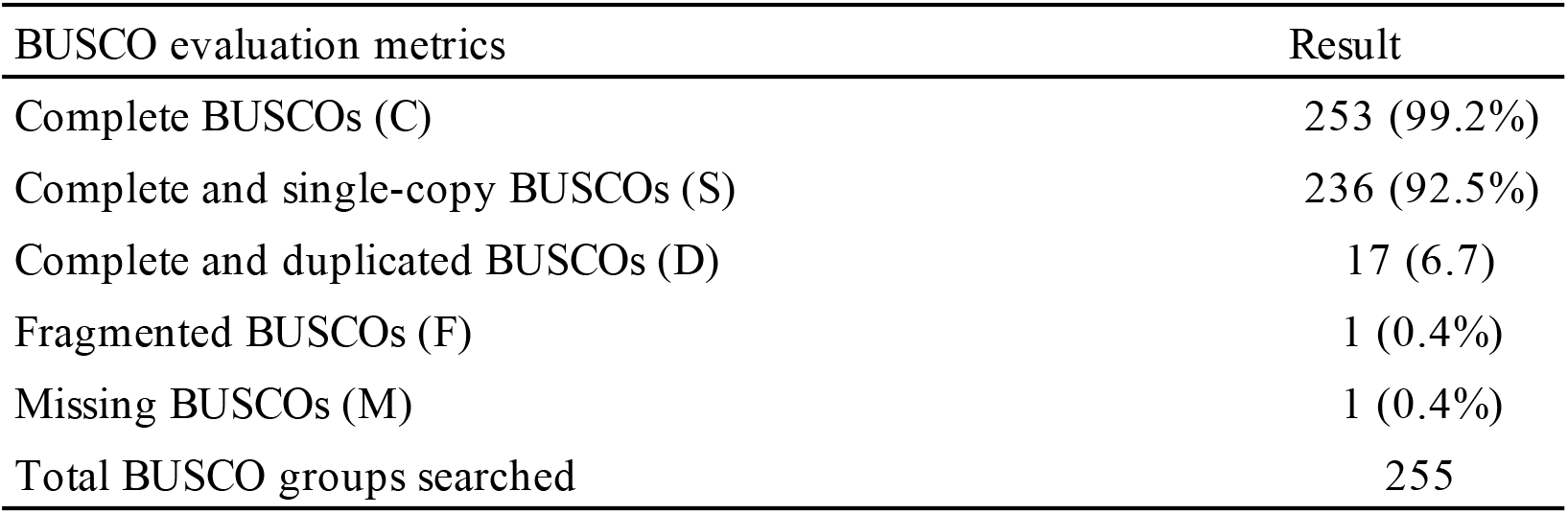
Transcript completeness from BUSCO analysis.

### 3.2 Time-course transcriptomic profiles during embryonic development under MPT conditions

We conducted the time-course gene expression analysis at MPT to investigate the embryonic gonadal transcriptome leading to the induction of functional testis differentiation in *M*. *reevesii*. We successfully extracted eight groups ("UUU", "UUE", "UEE", "UEU", "EUU", "EUE", "EEU", and "DDD") of transcripts with statistically up-regulated patterns during developmental stages 15-18 (Figure 1B; Table S1). Here, "UUU" and "UEE" indicate the up-up-up and up-equal (no change)-equal expression patterns in stages 15 to 16, 16 to 17, and 17 to 18, respectively. Previous studies have reported male-linked genes up-regulated at MPT in the red-eared slider turtle *T*. *scripta* (Czerwinski et al., 2016; Shoemaker et al., 2007), which include *Dmrt1*, *anti-Müllerian hormone* (*Amh*), and *SRY-box transcription factor 9* (*Sox9*). They are all conserved genes with known functions in male sexual differentiation in vertebrates including reptile species (Kohno et al., 2014; Morrish and Sinclair, 2002; Valenzuela et al., 2013). In *M*. *reevesii*, the expression level of *Dmrt1* was progressively up-regulated depending on the developmental stage (Figure 2) and was categorized into the "UEU" group (Figure 1B). In *T*. *scripta* (Czerwinski et al., 2016), differential expression of *Dmrt1* between males and females occurs at stage 16, and the expression level of *M. reevesii Dmrt1* at stage 16 at MPT is consistently higher than in stage 18 at FPT (Figure 2). These data are similar to the findings for a previous qPCR assay for *Dmrt1* in *M. reevesii* (Dong et al., 2020). *Sox9* exhibited a similar expression pattern to *Dmrt1* ("UEU" group) (Figure 2). The expression level of *Amh* gradually increased between stages 15-17 and then increased dramatically at stage 18 ("UUU" group) (Figure 2), which is consistent with the *Amh* expression pattern in *T*. *scripta* (Czerwinski et al., 2016). In a previous study of *M. reevesii* (Dong et al., 2020), distinct sexual differences appeared after stage 21, which differs somewhat from the results of this study, but they are consistent in the context that these differences appeared in the late TSP stage. Thus, transcriptional regulation underlying male sexual differentiation is conserved between *M. reevesii* and *T*. *scripta*.

**Figure 2.**
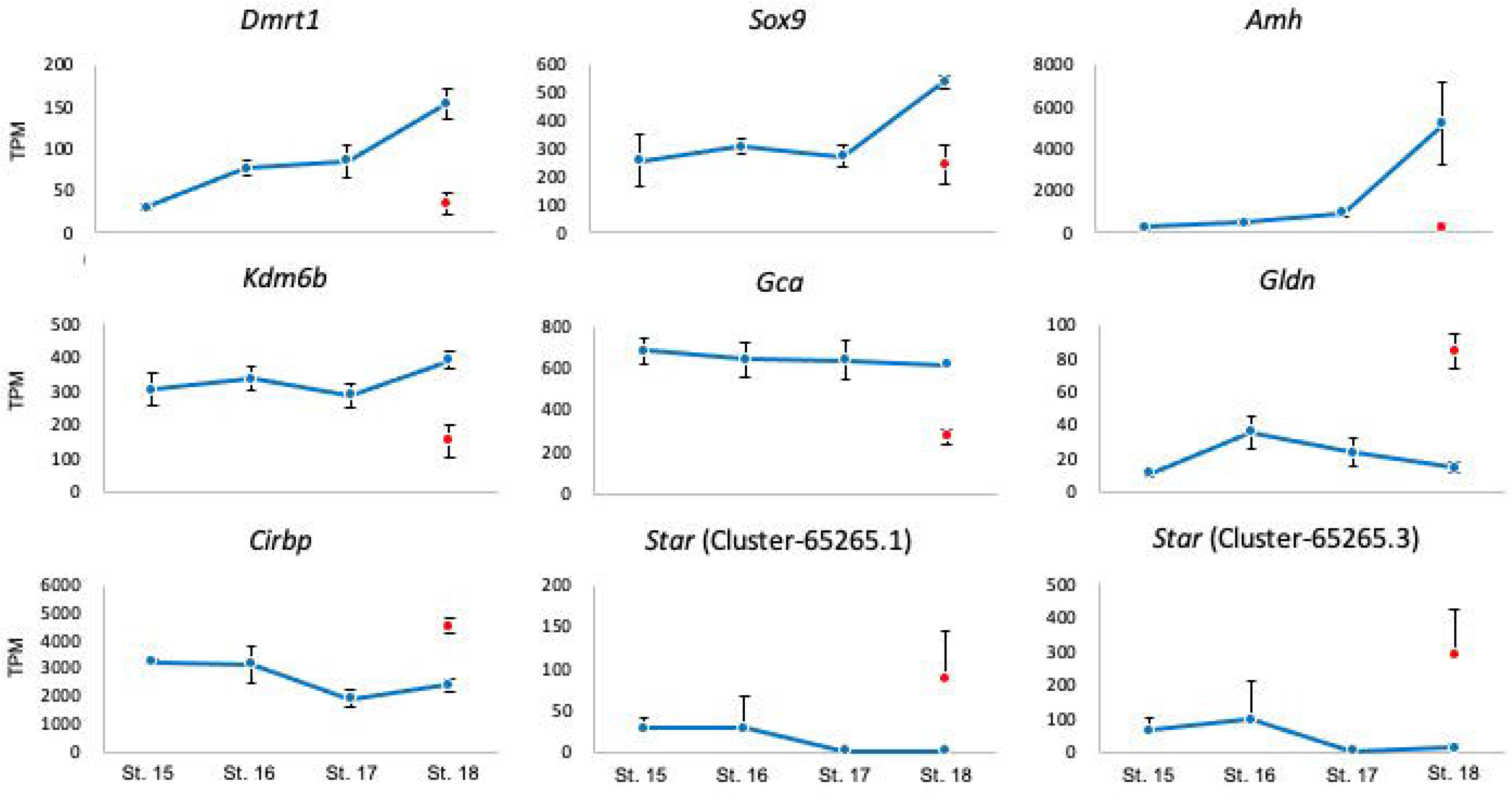
Gonadal expression profiles of selected differentially expressed genes between MPT and FPT at stage 18of *M*. *reevesii*. MPT profiles cover stages 15-18 (blue dots) whereas the red point indicates expression at FPT. Bars indicate standard deviations. TPM indicates the transcripts per million. Abbreviations of gene names are as follows: *Dmrt1* (*doublesex and mab-3 related transcription factor 1*), *Sox9* (*SRY-box transcription factor 9*), *Amh* (*anti-Müllerian hormone*), *Kdm6b* (*Lysine (K)-specific demethylase 6B*), *Gca* (*grancalcin*), *Gldn* (*gliomedin*), *Cirbp* (*cold-inducible RNA binding protein*), and *Star* (*steroidogenic acute regulatory protein*).

To provide an overview of the molecular events following the developmental time-course at MPT, we carried out GO and KEGG enrichment analyses using genes in the "UUU" and "DDD (down-down-down expression pattern)" groups. In the "UUU" group, the metabolic pathway terms (KEGG: mmu01100; GO: lipid metabolic process) and developmental pathway-related GO terms (e.g., positive regulation of cell migration and animal organ morphogenesis) were enriched (Table 2). On the other hand, the cell cycle-related GO terms (e.g., cell cycle and cell division), RNA processing-related GO terms (e.g., mRNA processing and RNA splicing), and apoptosis-related GO term (apoptotic process) were enriched in "DDD" group (Table 2). In recent years, differential alternative splicing has been found to occur in several genes depending on incubation temperature. Intron retention, the insertion of introns in mRNA sequences after the transcription, could cause translation termination due to the frame shifts or inserted termination codons, resulting in nonfunctional protein production. In *T*. *scripta*, alternative splicing at MPT has led to the retention of the 19th intron of *Lysine (K)-specific demethylase 6B* (*Kdm6b*) and the 15th intron of *jumonji AT rich interactive domain 2* (*Jarid2*) mRNA resulting in nonfunctional transcripts (Deveson et al., 2017). Moreover, an alternative splicing variant of the *Dmrt1* in which exons 2 and 3 were skipped (Dmrt1 ΔEx2Ex3) was found in the painted turtle *C*. *picta* (Mizoguchi and Valenzuela, 2020). Here, our RNA-seq data demonstrated that expression profiles of several genes with RNA processing-related GO terms were lowered depending on developmental stage progression, implying that alternative splicing is involved in sex differentiation-related genes such as *Dmrt1* in *M. reevesii*. Additionally, the results showed that genes related to the metabolic pathway and organ morphogenesis are highly expressed at stages 15-18 at MPT. These genes include *prostaglandin-endoperoxide synthase 2* (*Ptgs2*), *vanin 1* (*vnn1*), *regucalcin* (*Rgn*), *glyoxylate reductase/hydroxypyruvate reductase* (*Grhpr*), *3-hydroxybutyrate dehydrogenase type 1* (*Bdh1*), *carnosine dipeptidase 1* (*cndp1*). Of these genes, *vnn1*, a family of pantetheinase enzymes, is expressed in male-developing gonads in mice (Bowles et al., 2000; Wilson et al., 2005). These results clearly indicate that the coordination of multiple processes leads to the transformation of undifferentiated gonads into testes.

**Table 2.**
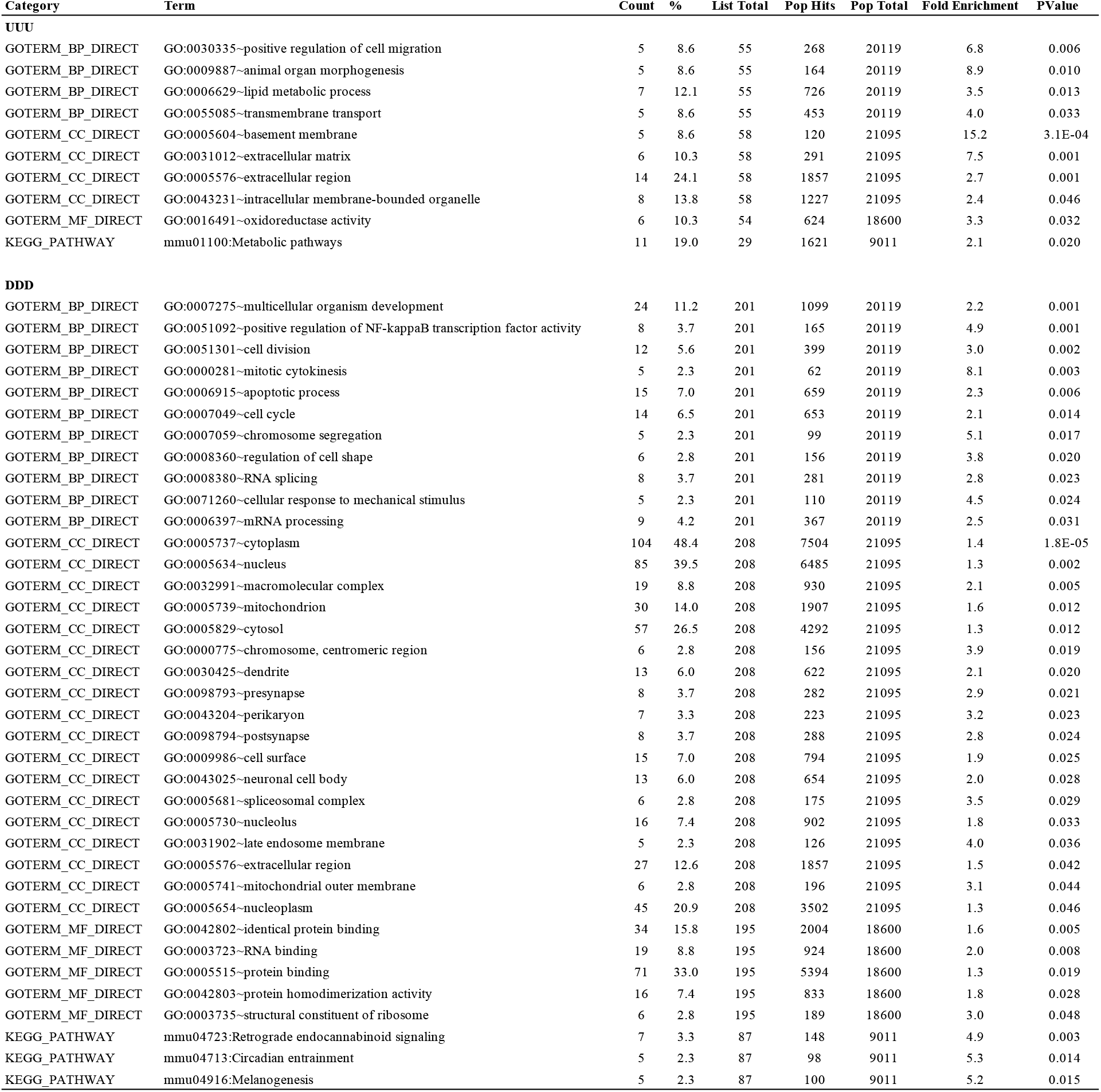
List of gene ontology and KEGG pathway enriched in UUU and DDD categories

### 3.3 Differentially expressed genes between MPT and FPT conditions

Among 113,460 constructed transcripts, 66 and 41 genes were screened as DEGs (false discovery rate < 0.05) in the MPT and FPT at stage 18 of *M. reevesii*, respectively (only annotated DEGs indicated in Table 3). Principal component analysis of the entire gene expression dataset indicated that the transcriptome profiles of MPT and FPT at stage 18 were clearly separated (Figure S1A), including the MPT-biased genes, *Dmrt1*, *Sox9*, and *Amh* (Figure 2). In addition to transcriptional regulation, epigenetic marks, such as DNA methylation and histone modifications of known regulatory genes of gonadal sexual differentiation, have been shown to differ between MPT and FPT in TSD species (Dong et al., 2020; Matsumoto et al., 2013, 2016; Parrott et al., 2014; Piferrer, 2013; Radhakrishnan et al., 2017). Although these reports were not clear on whether differences in epigenetic status are a cause or a consequence of sexual development in TSD species, recently two factors, *Kdm6b* (histone demethylase) and *signal transducer and activator of transcription 3* (*Stat3*), have been shown to play central roles in epigenetic regulation in reptiles (Ge et al., 2018; Weber et al., 2020). Stat3 is phosphorylated at the warmer temperature (FPT), it binds the Kdm6b locus, and represses *Kdm6b* transcription, resulting in an inhibition of the male pathway. Therefore, the *Stat3* expression pattern itself does not change significantly between MPT and FPT during TSP in *T. scripta* (Weber et al., 2020). Our transcriptome data showed that the expression pattern of *Stat3* did not differ between MPT and FPT at stage 18 (average transcript per million (TPM): ca 400). By contrast, *Kdm6b* exhibited the dimorphic expression pattern at MPT and FPT (Figure 2), but an elevated expression pattern was not observed during MPT development, unlike that in *T. scripta* (Ge et al., 2018).

**Table 3.**
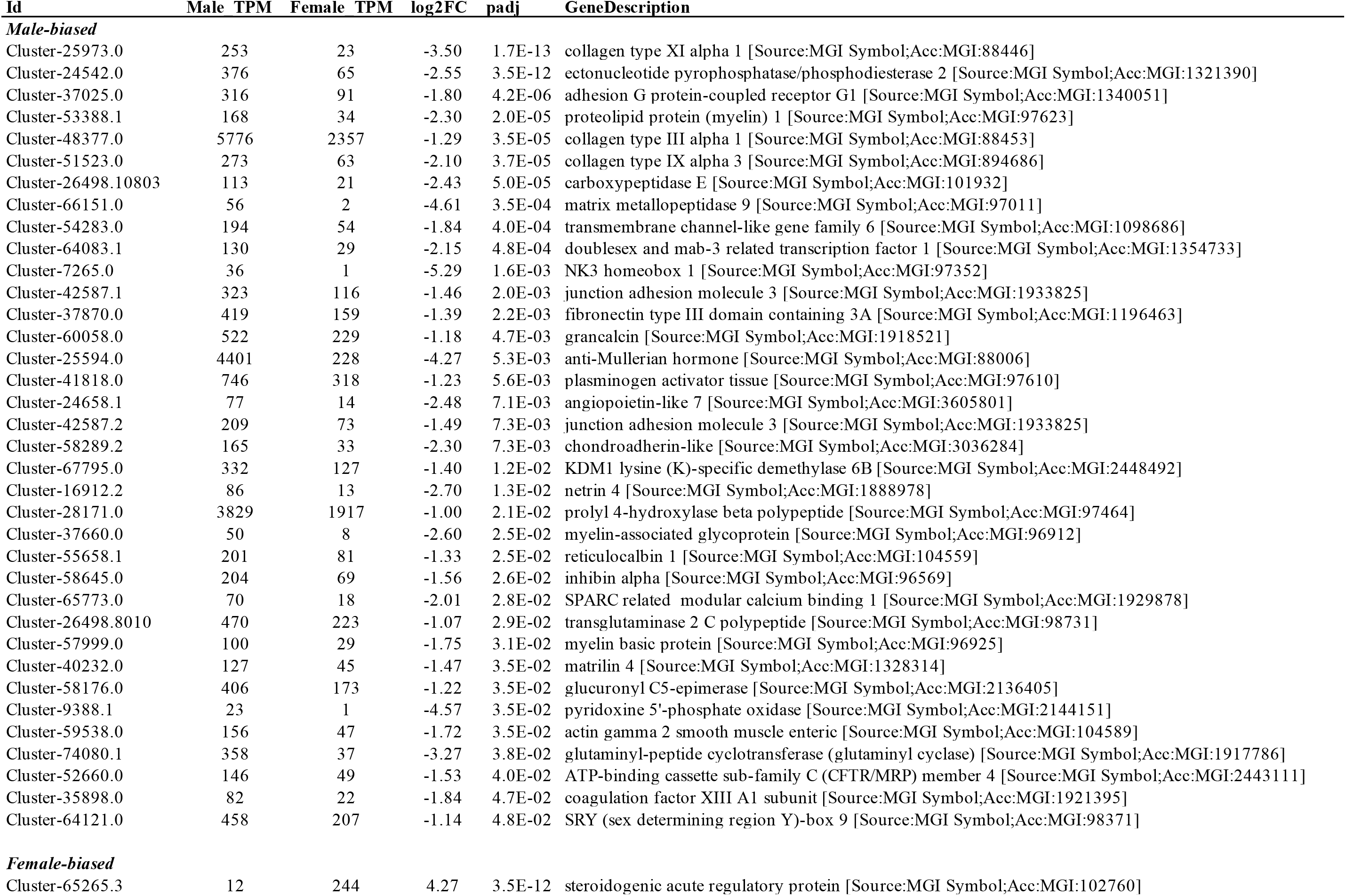
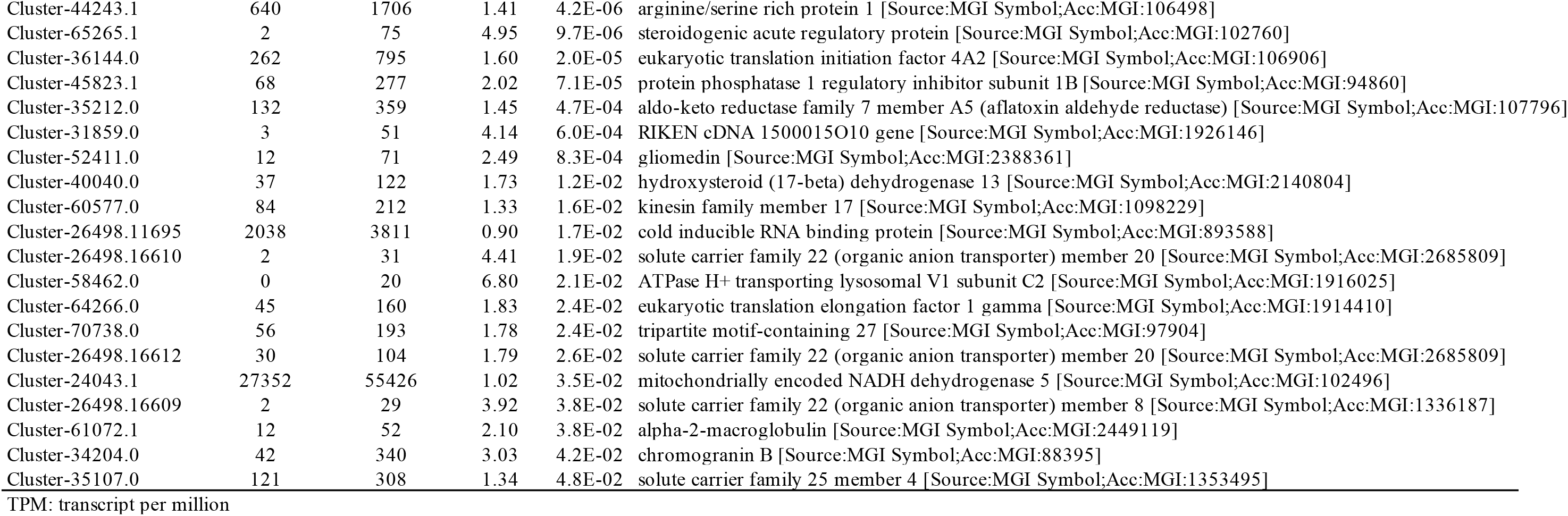
List of male- and female-biased differential expressed transcripts

Another factor known to often be associated with temperature during TSP is *grancalcin* (*Gca*). This gene encodes a calcium-binding protein that is commonly found in neutrophils and macrophages in mice (Roes et al., 2003) and is associated with ovarian development in the central bearded dragon *P*. *vitticeps,* a GSD reptile (Whiteley et al., 2021). Expression of *Gca*, however, was significantly higher in MPT than in FPT at stage 18 in *M. reevesii* (Figure 2). On the other hand, FPT-biased DEGs contained *gliomedin* (*Gldn*), *cold inducible RNA binding protein* (*Cirbp*), and two transcripts of *steroidogenic acute regulatory protein* (*Star*) (Figure 2). *Gldn* encodes a protein required for the formation of the nodes of Ranvier and the development of the human peripheral nervous system (Wambach et al., 2017). In *T. scripta*, *Gldn* is specifically expressed in FPT from the early stages of TSP increasing during FPT incubation (Czerwinski et al., 2016), although its function remains unknown. *Cirbp* is known to regulate temperature-dependent cellular processes via translational repression and mRNA stabilization (De Leeuw et al., 2007; Liu et al., 2013; Xia et al., 2012; Yang et al., 2006), and is suggested to be involved in sexual fate decision in the common snapping turtle, *Chelydra serpentina* (Schroeder et al., 2016). *Star* is important for steroid biosynthesis and the female-biased gene expression is plausibly associated with estrogen production. On the other hand, *forkhead box L2* (*Foxl2*) and *cytochrome P450 family 19 subfamily a member 1* (*Cyp19a1*; also known as *aromatase*), which are direct effectors for female sex differentiation genes widely conserved in vertebrates, were not detected in FPT at stage 18 in *M. reevesii*, and this is consistent with the previous qPCR-based analysis (Dong et al., 2020). Moreover, their expression increased late in the gonadal differentiation period in *T*. *scripta* (Czerwinski et al., 2016) and *A*. *mississippiensis* (Yatsu et al., 2016). Hence these genes may be upregulated after stage 19 in *M. reevesii*. The repertoire of DEGs indicates that the gene cascade underlying the early-to-middle period of TSP in *M. reevesii* is comparable with other TSD species.

### 3.4 Comparison of common DEGs between M. reevesii and A. mississippiensis

We compared the gonadal transcriptome at MPT and FPT of *M*. *reevesii* and *A*. *mississippiensis* to better understand the common transcripts between turtles and alligators. We re-analyzed our previous alligator RNA-seq data using only stages 22–3 at MPT (33.5[) and FPT (30[) (Yatsu et al., 2016). Among 126,264 constructed transcripts, 170 and 80 genes were screened as DEGs (false discovery rate < 0.05) in the MPT and FPT at stages 22-23 of *A*. *mississippiensis*, respectively. Principal component analysis of the entire gene expression dataset indicated that the transcriptome profiles of MPT and FPT in stages 22-23 were clearly separated (Figure S1B). A notable point of this analysis is that the temperature ranges of MPT and FPT are opposite to on another for *M. reevesii* and *A*. *mississippiensis* (for *M. reevesii* FPT is at high temperatures and MPT at low temperatures, whereas for *A*. *mississippiensis* MPT is at high temperatures and FPT at low temperatures). This makes it possible to extract male- or female-linked genes and high- or low-temperature-responsive genes that are common/different to both by directly comparing the expression patterns of functionally-homologous genes. Based on this comparative analysis, two genes, *matrilin 4* (*Mtn4*) and *inhibin alpha* (*Inha*), were found as common male-biased genes, in addition to well-conserved male sexual differentiation genes, such as *Amh*, and *Sox9* (Figure 3). Interestingly, expression levels of both *Mtn4* and *Inha* increased from stage 17 at MPT in *M. reevesii* (Figure 4). Inha forms dimers with the Inhb subunit to form inhibin and regulates a broad range of reproductive processes including steroidogenesis and germ cell maturation (Barakat et al., 2012; Itman et al., 2006). Inhibin is a member of the transforming growth factor-β (TGFβ) family member as Amh is. However, the involvement of inhibin in gonadal differentiation remains unclear.

**Figure 3.**
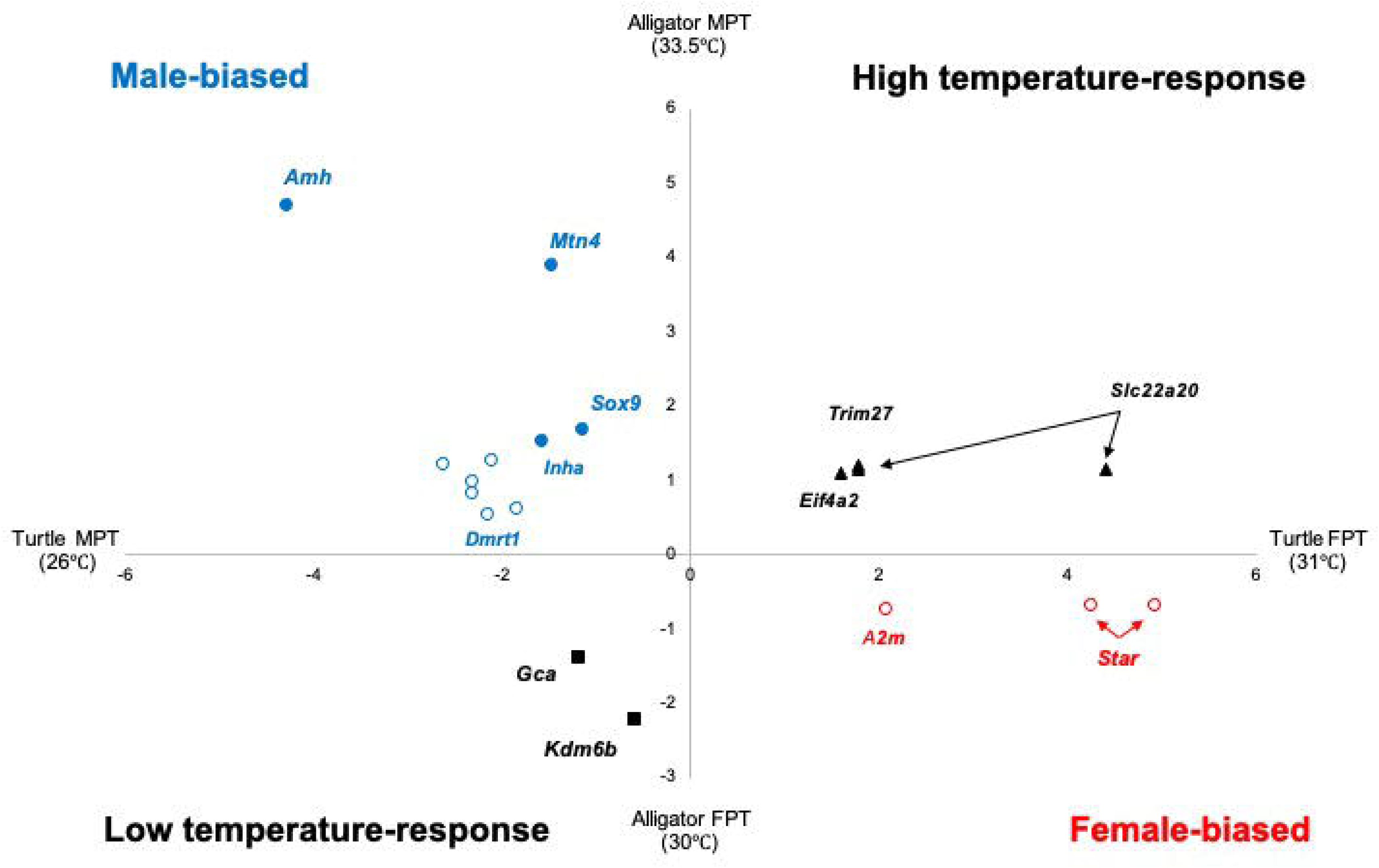
Gonadal expression profiles of common differentially expressed genes between MPT and FPT of the Chinese pond turtle *M*. *reevesii* (stage 18) and American alligator *A. mississippiensis* (stage 22-23). X-axis and y-axis indicate the log_2_FC (fold change). The blue points are categorized as "male-biased" genes with FDR &lt; 0.05 for both turtles and alligators. The blue circles are also categorized as "male-biased" genes with FDR &lt; 0.05 for turtles and FDR > 0.05/ log_2_FC > |0.3| for alligators. The red circles are categorized as "female-biased" genes with FDR &lt; 0.05 for turtles and FDR > 0.05/ log_2_FC > |0.3| for alligators. Triangles and squares are genes (FDR &lt; 0.05 for both turtles and alligators) that indicate "high-temperature response" and "low-temperature response", respectively. Abbreviations of gene names are as follows: Mtn4 (*matrilin 4*), Inha (*inhibin alpha*), A2m (*Alpha-2-macroglobulin*), *Trim27* (*tripartite motif-containing 27*), *Eif4a2* (*eukaryotic translation initiation factor 4A2*), and *Slc22a20* (*solute carrier family 22 member 20*).

**Figure 4.**
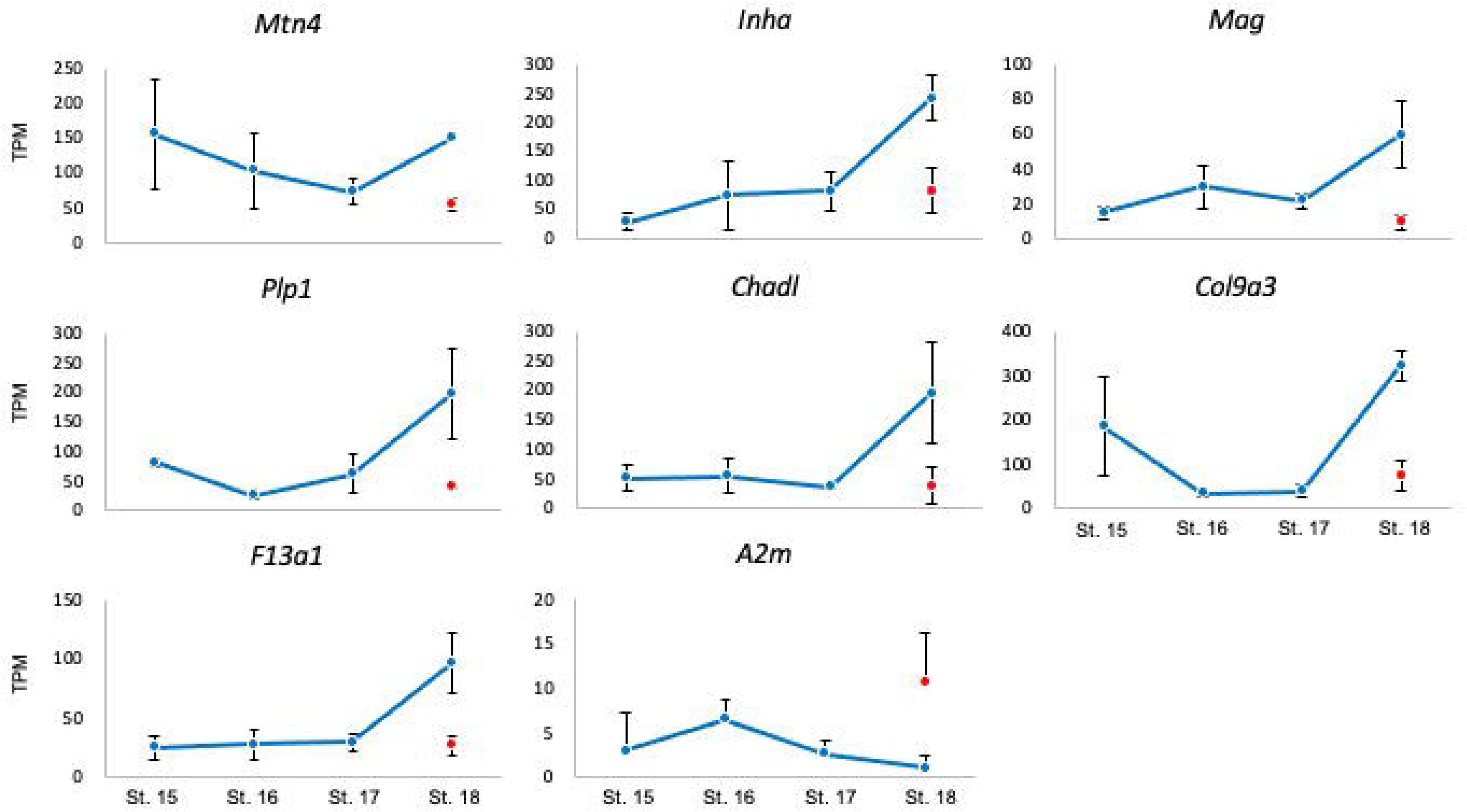
Gonadal expression profiles of selected differentially expressed genes between MPT and FPT in stage 18of *M*. *reevesii*. MPT profiles cover stages 15-18 (blue) and FPT indicates the red point. Bars indicate standard deviations. TPM indicates the transcript per million. Abbreviations of gene names are as follows: *Mag* (*myelin-associated glycoprotein*), *Plp1* (*proteolipid protein 1*), *Chadl* (*chondroadherin-like*), *Col9a3* (*collagen type IX alpha 3*), and *F13a1* (*coagulation factor XIII A1 subunit*

*Myelin-associated glycoprotein* (*Mag*), *proteolipid protein (myelin) 1* (*Plp1*), *Chondroadherin-like* (*Chadl*), *Collagen type I*X *alpha 3* (*Col9a3*), and *Coagulation factor XIII A1* (*F13a1*), three transcripts of *Alpha-2-macroglobulin* (*A2m*) and two transcripts of *Star* were extracted as common male- and female-biased genes (genes with FDR < 0.05 for turtles and FDR > 0.05/ log_2_FC > |0.3| for alligators) (Figures 3, 4). Re-analysis of our previous alligator RNA-seq data demonstrated that two *Star* transcripts were predicted as commonly female-biased (Figure 3), indicating that Star might play an important role in steroidogenesis, probably estrogen production, in females in TSD reptiles. Thus, the male-biased and female-biased genes shared by alligators and turtles were nervous system (myelin)-related, collagen metabolism-related, blood metabolism-related, and steroidogenesis-related factors. Although these factors have not been the focus of previous TSD studies, it appears that they may be related to gonadogenesis in the TSP across TSD species. *Col9a3* is expressed specifically in the mouse testis after 11.5 days postcoitum and may contribute to testicular tubule morphogenesis throughout vertebrates (McClive and Snclair, 2003; Perera et al., 2001).

Although *Kdm6b* was identified as a key regulator for testicular differentiation in turtles, it was expressed at lower temperatures (FPT for alligator TSD) (Figure 3). Thus, *Kdm6b*-mediates not only gonadal differentiation but also other phenomena related to low-temperature response in alligators. Moreover, this analysis provided new insight into *Gca.* Comparison between *M. reevesii* and *A*. *mississippiensis* demonstrated that *Gca* was categorized as a low-temperature responsive gene (Figure 3). This is the opposite case for the temperature response of the central bearded dragon *P. vitticeps*, where the *Gca* expression is decreased at high-temperature incubation in this GSD lizard (Whiteley et al., 2021). Thus, *Gca* may have a different response temperature range for each species, but at least in the gonad during TSP, Gca may have a conserved function across species. This comparison of the transcriptome profiles between *M. reevesii* and *A*. *mississippiensis* TSPs indicates that the gene cascade for sexual differentiation is consistently activated earlier in males than in females.

## 4. Conclusion

Here, we conducted a time-series gonadal transcriptome analysis of the Chinese three-keeled pond turtle *M*. *reevesii* during TSP under MPT conditions, and demonstrate expression surges of well-conserved male sexual differentiating genes, such as *Amh*, *Sox9*, and *Dmrt1*, observed after stage 17. Furthermore, a comparison of MPT and FPT-biased genes in alligator embryonic stages 22-23, which correspond to stage 18 of the late TSP in *M*. *reevesii*, has resulted in the identification of new male- and female-biased genes. Notably, *Kdm6b*, known as a male-sex determiner in the turtle *T*. *scripta*, is highly expressed in FPT in alligators, suggesting that Kdm6b mediates not only gonadal differentiation but also other phenomena related to low-temperature response in alligators. Overall, our study provided new insights into molecular clues involved in the TSD of reptiles.

## Acknowledgments

We would like to thank Dr. Ryohei Yatsu for the sample collection and for establishing the experimental design. We also thank Drs. Kaori Miyaoku (Osaka University), Genki Yamagishi (Tokyo University of Science), Albert Zhou (University of Birmingham, UK), and Hiroyo Nishide (National Institute for Basic Biology, Japan) for their technical support for data analysis. This work was supported by Grants-in-Aid for Scientific Research from the Ministry of Education, Culture, Sports, Science and Technology of Japan (21H02522, 17H06432 to S.M.), NIBB Collaborative Research Program (19-405, 20-415, 21-310 to S.M.), Ministry of the Environment, Japan (T.I.), and computational resources were provided by the Data Integration and Analysis Facility, National Institute for Basic Biology.

## Data availability

The raw-read datasets were deposited in DDBJ under the accession number: DRA016380. Other data will be made available upon reasonable request.

---

**Figure S1.**
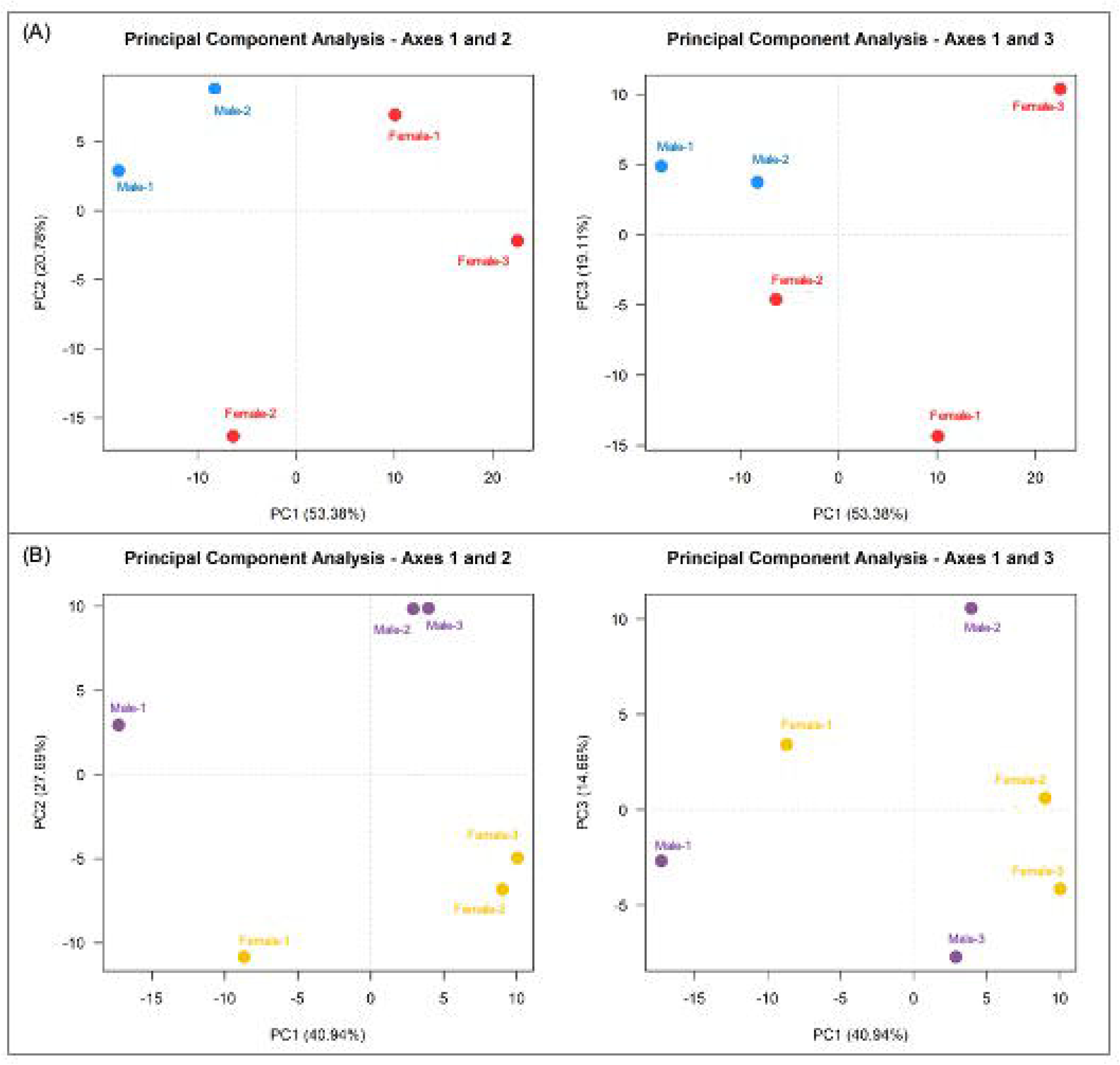
Principal component analysis (PCA) of the transcriptome analysis data obtained from the gonads under MPT and FPT at stage 18 *M. reevesii* (A) and at stage 22-23 *A. mississippiensis* (B).

**Table S1.**
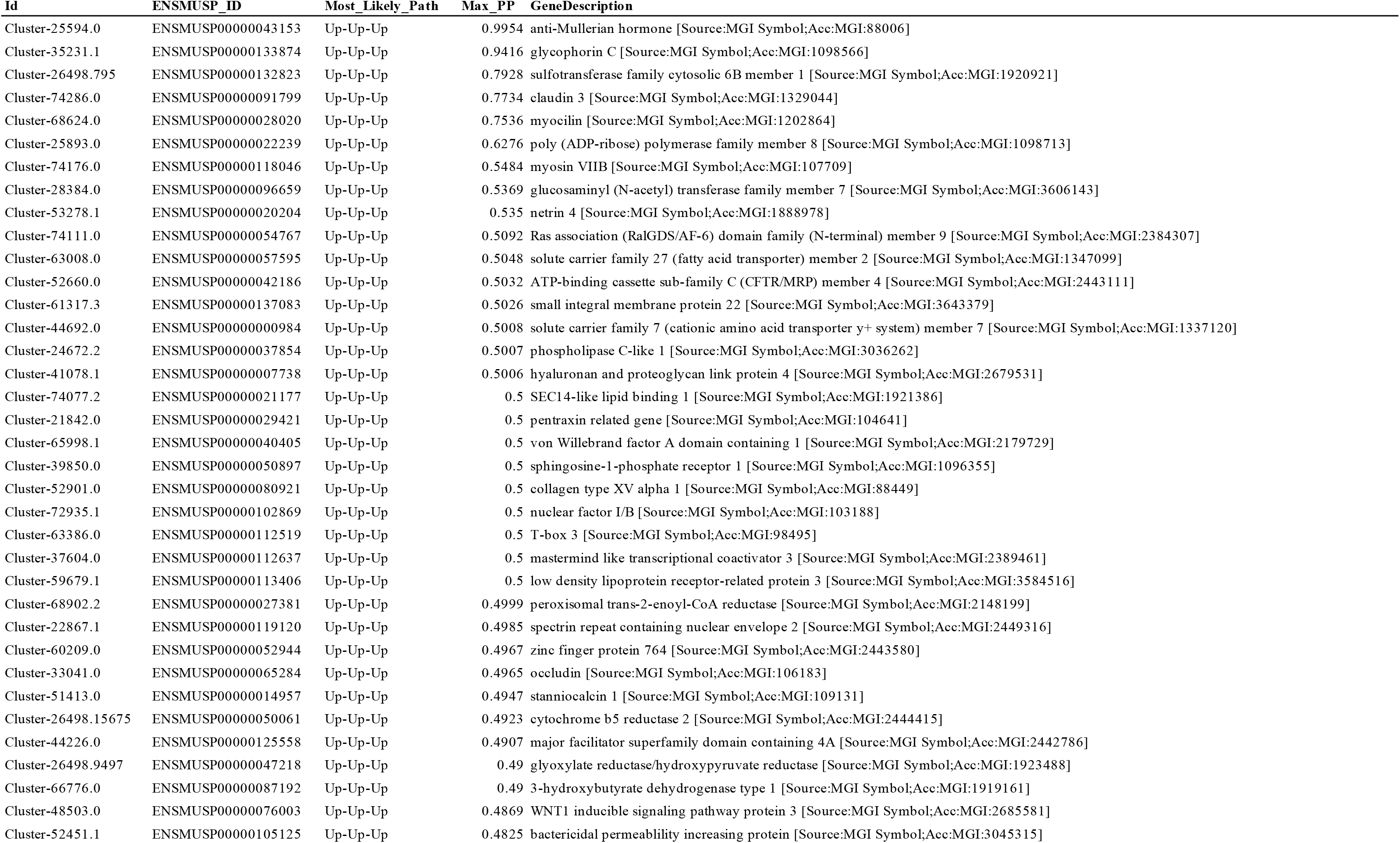
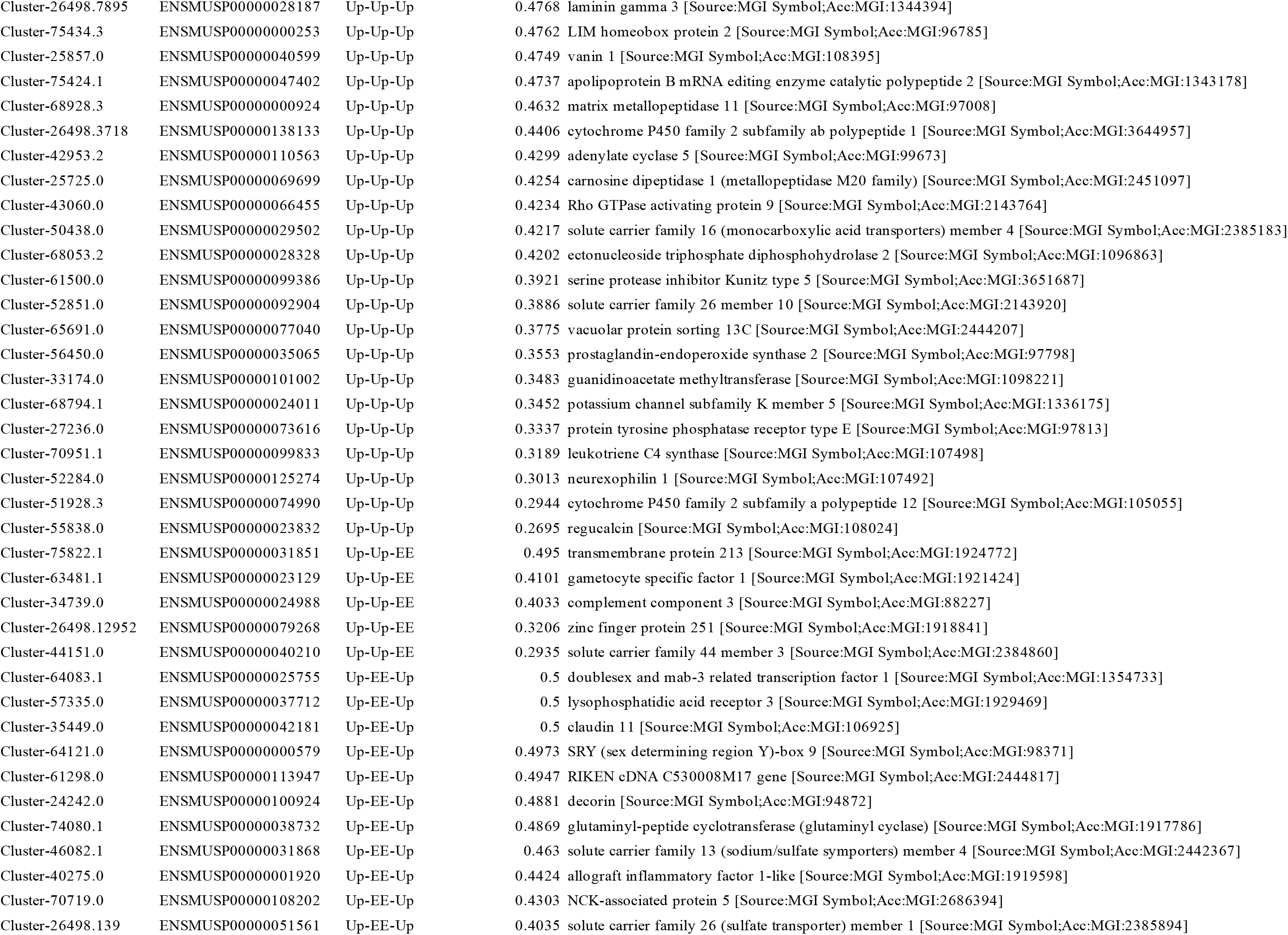
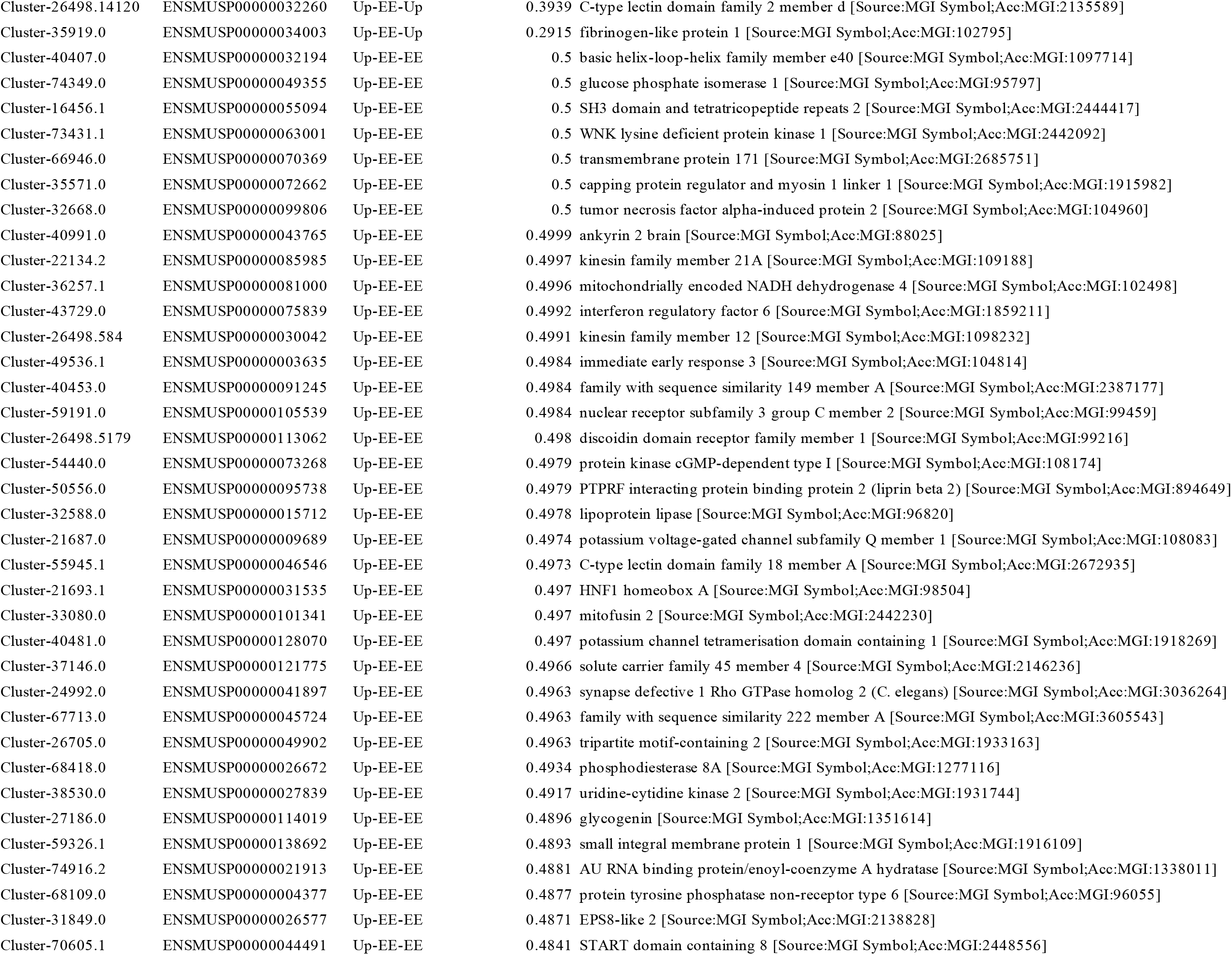
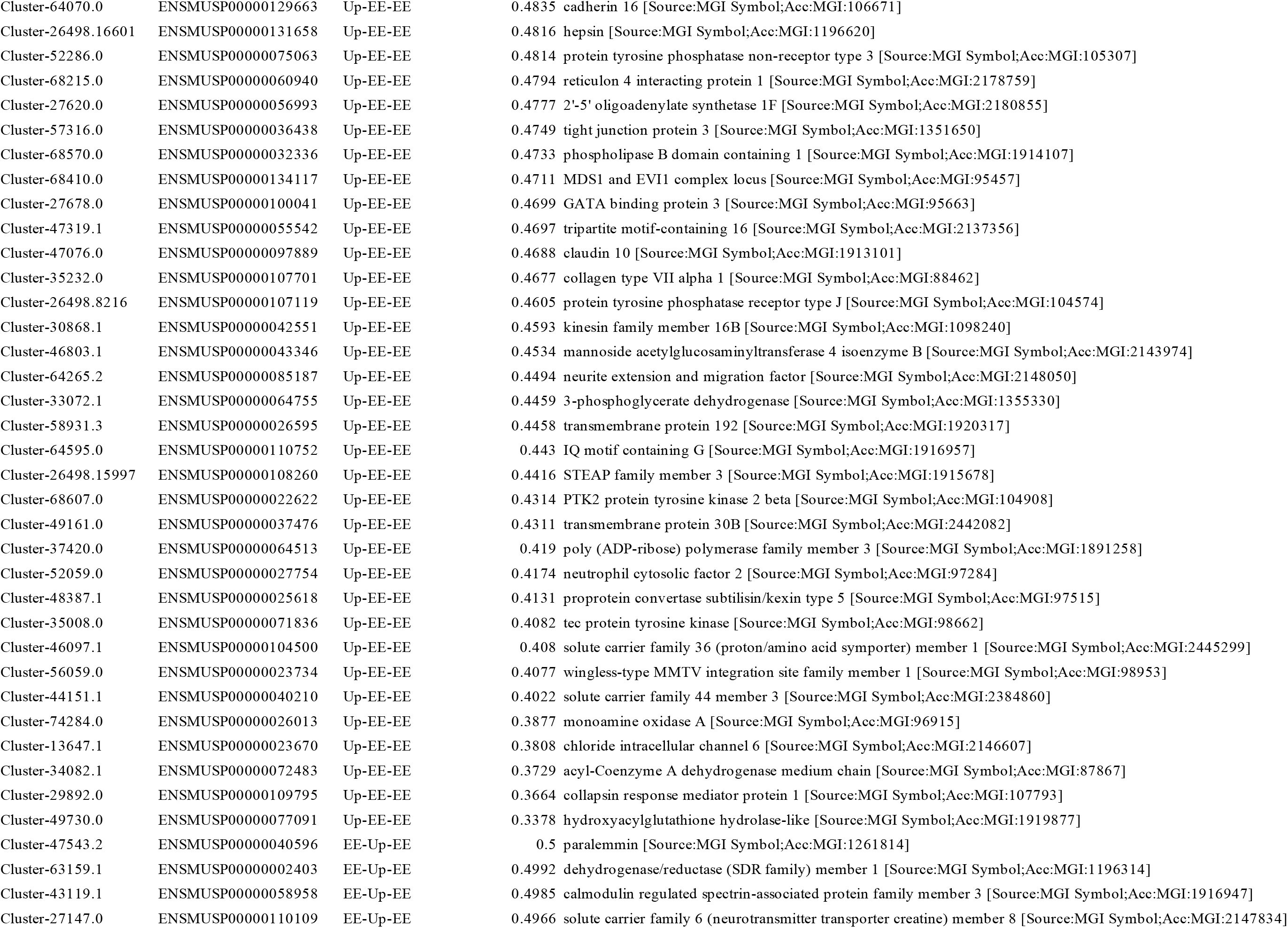
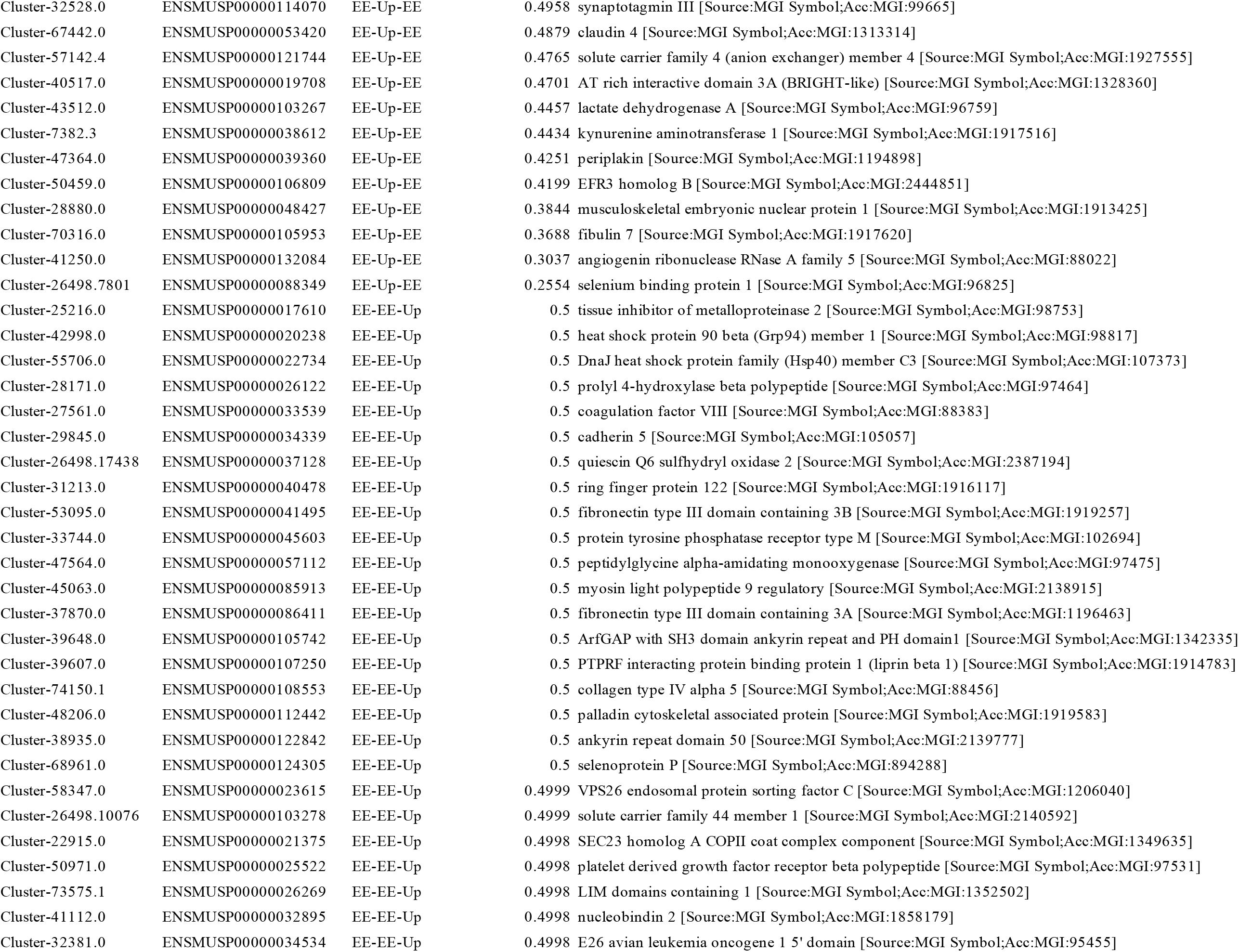
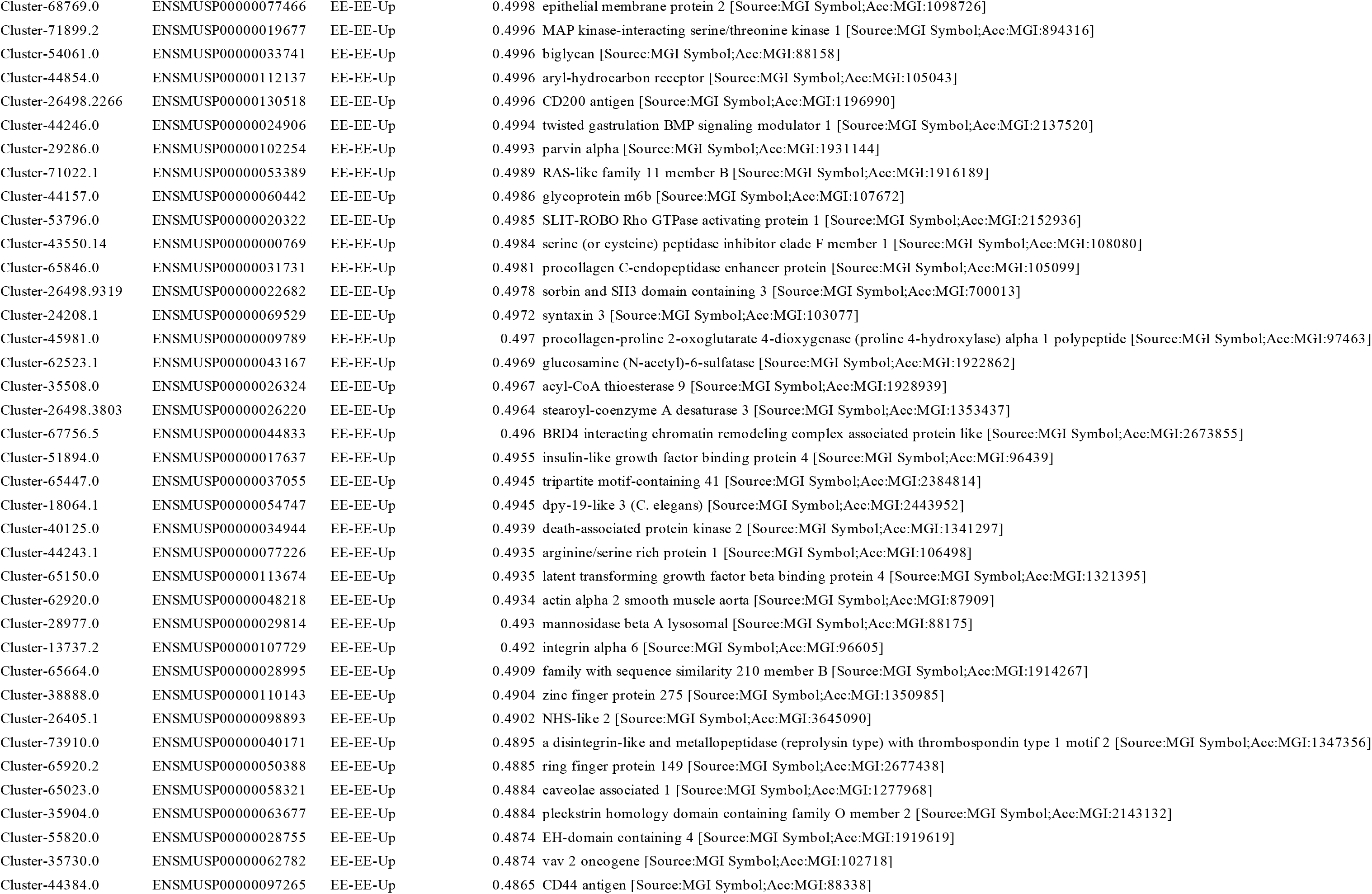
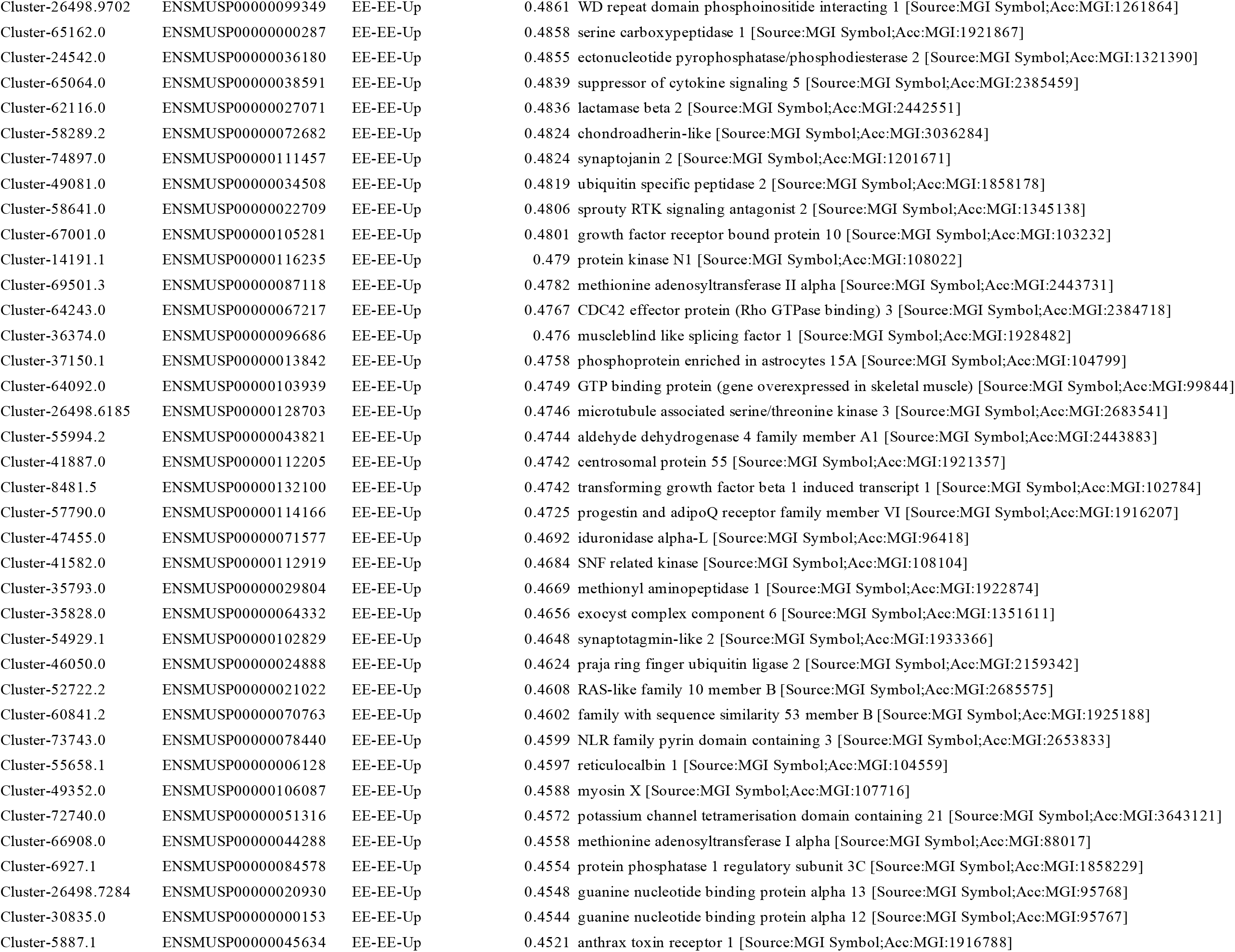
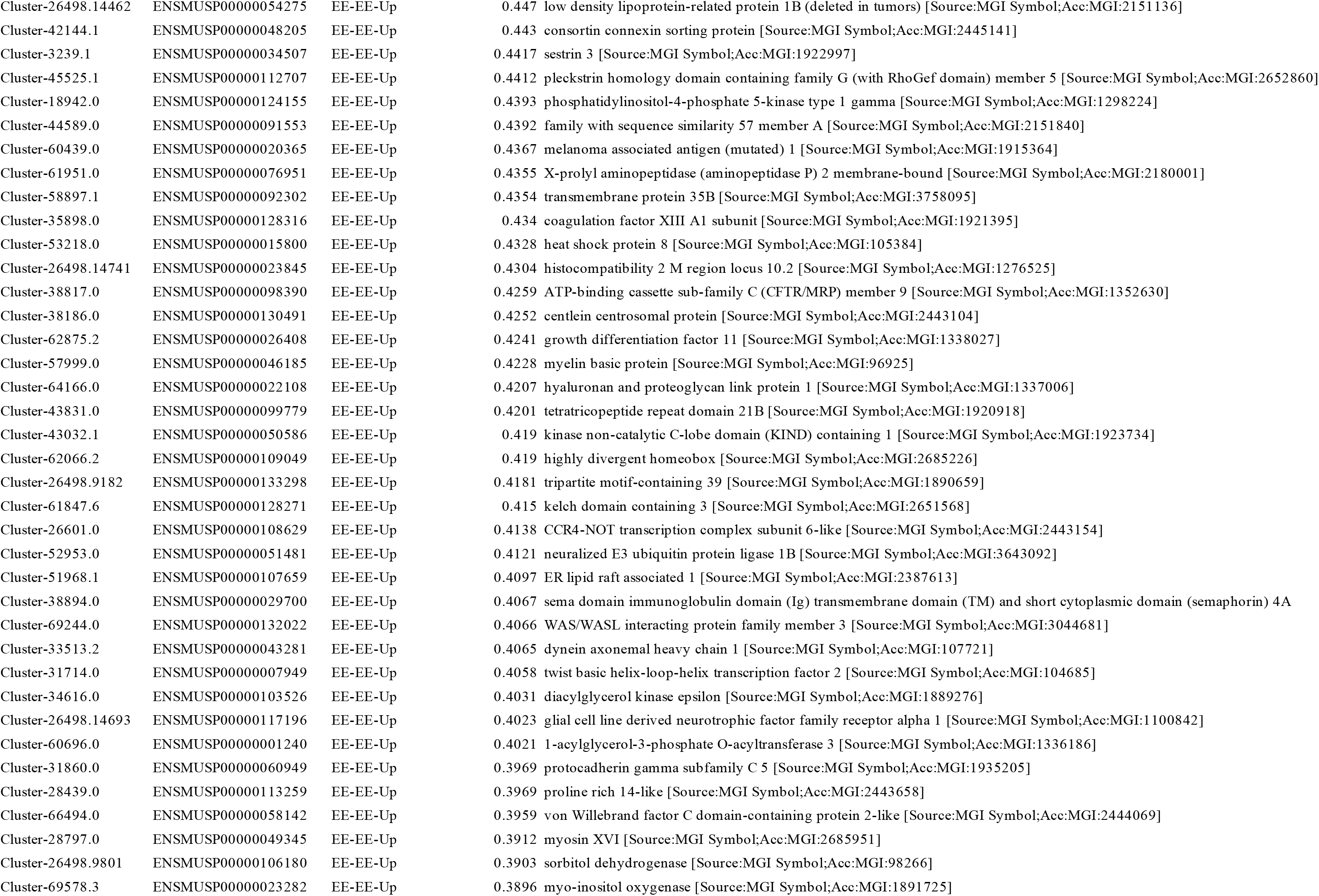
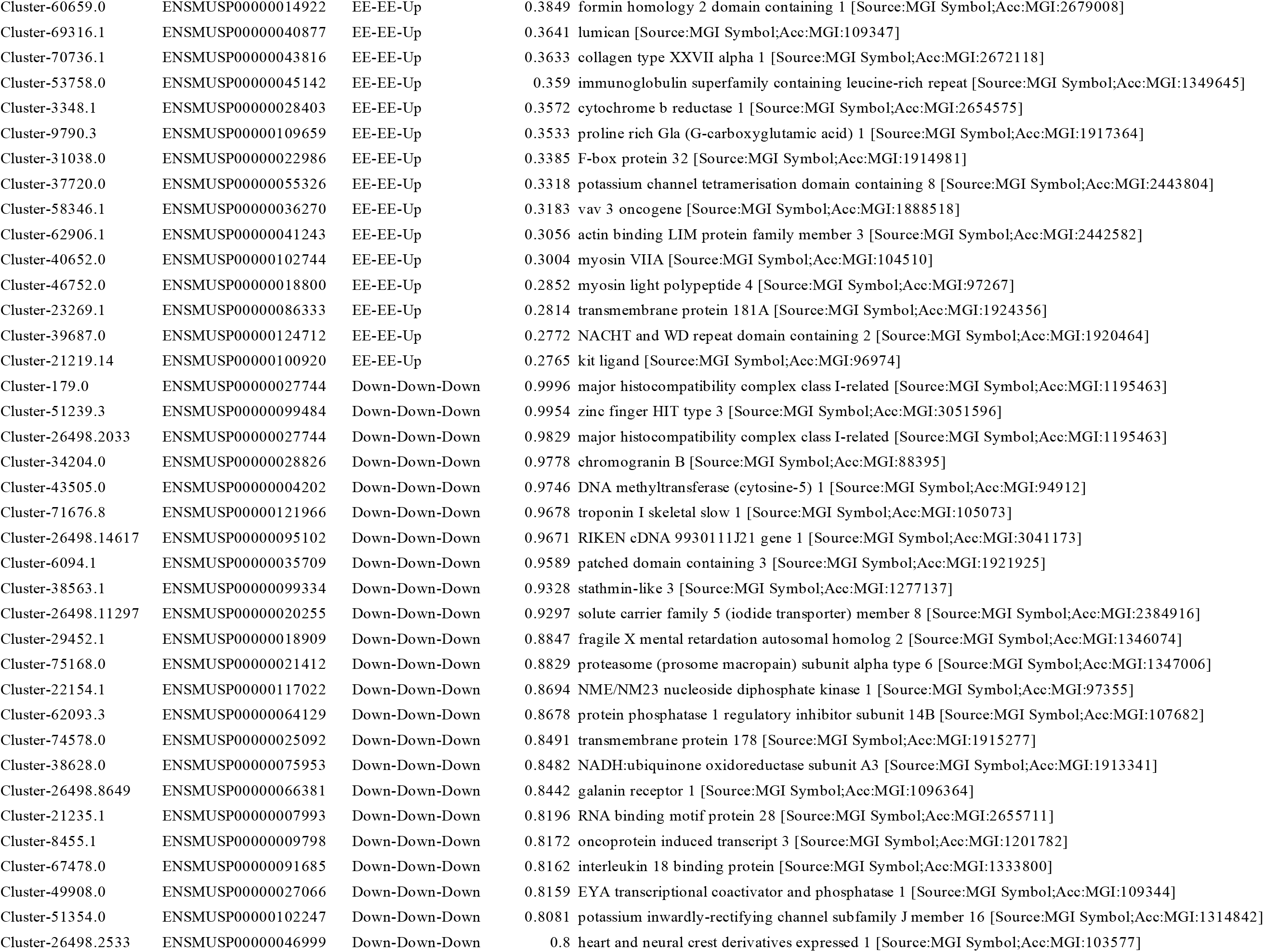
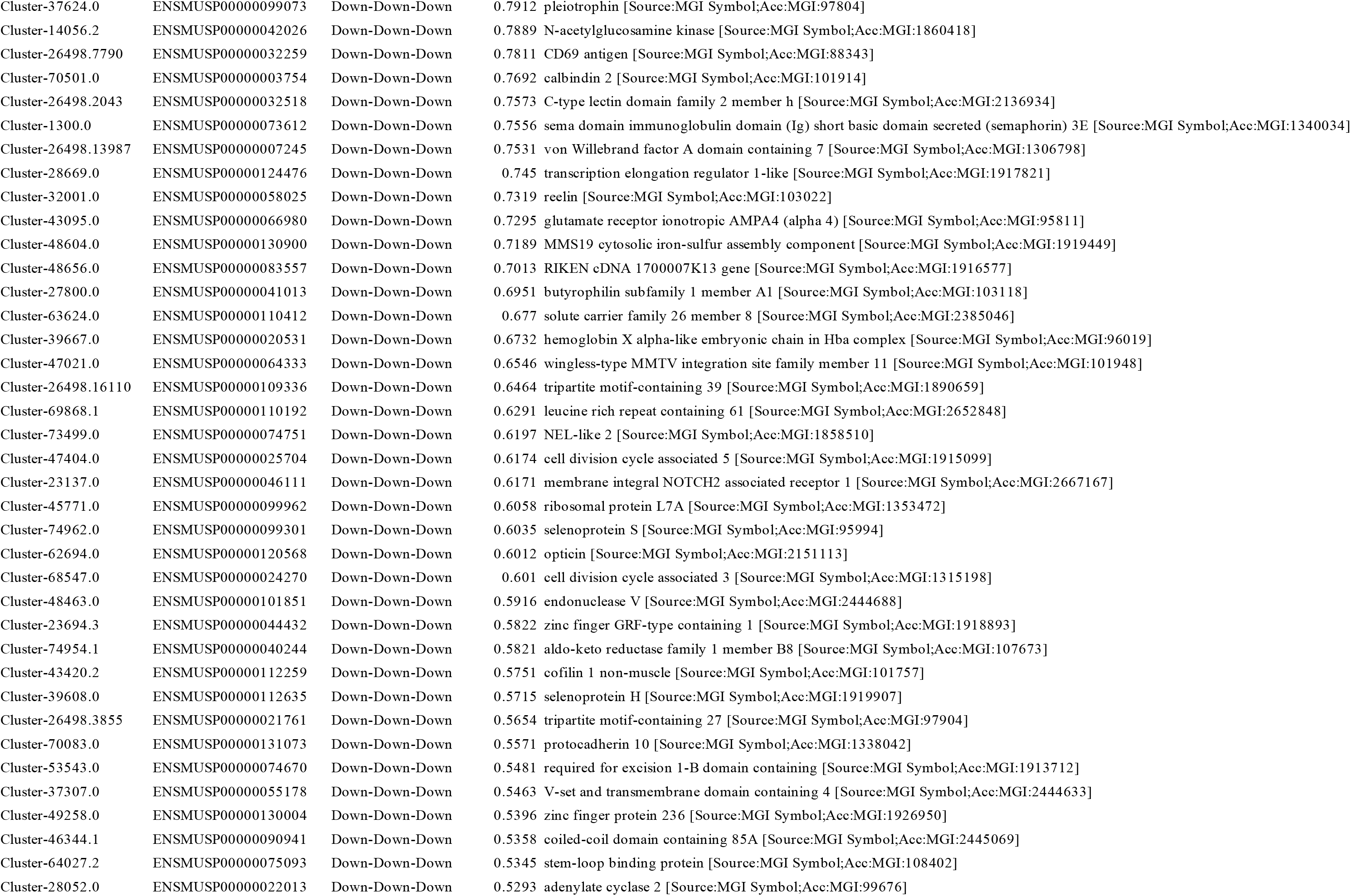
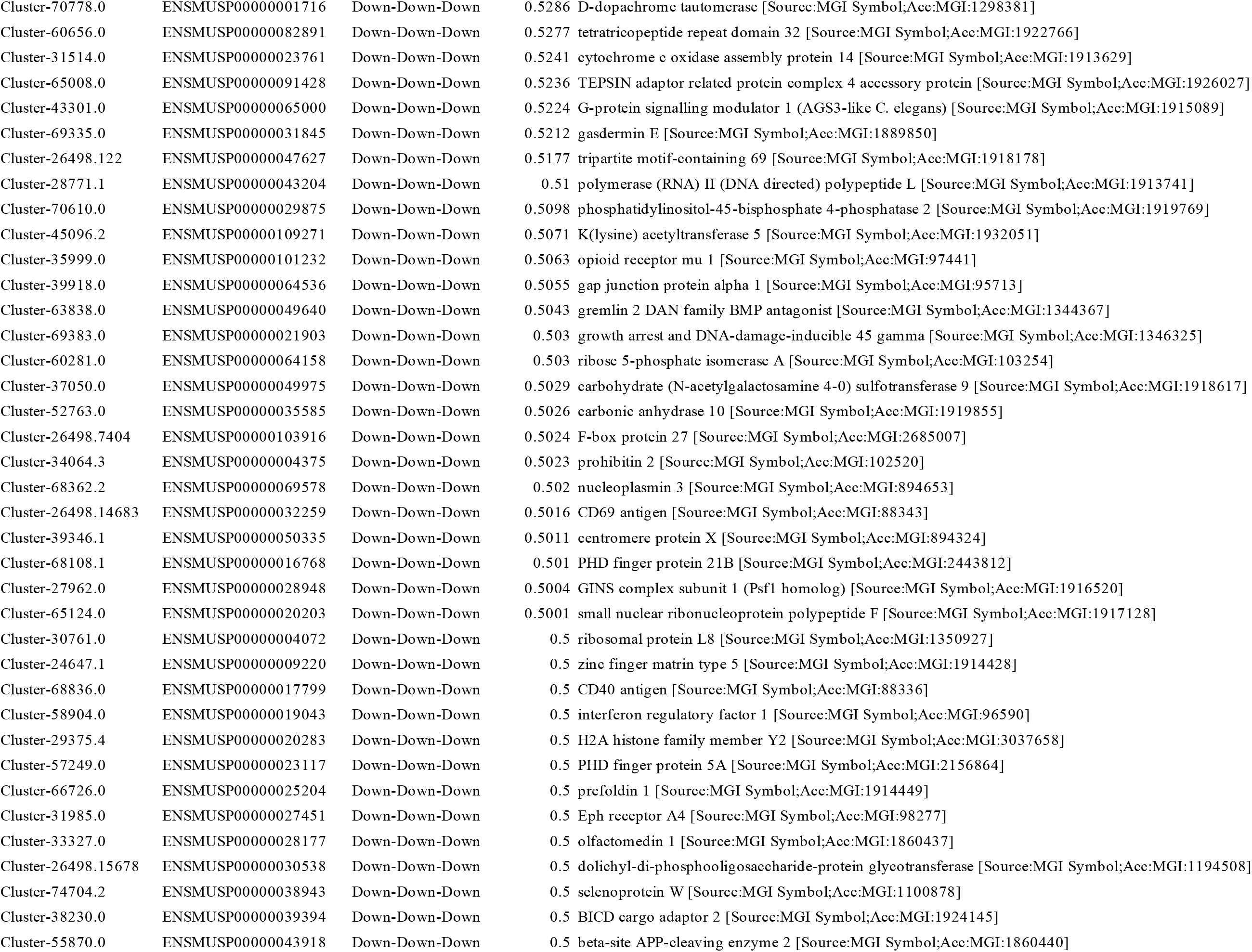
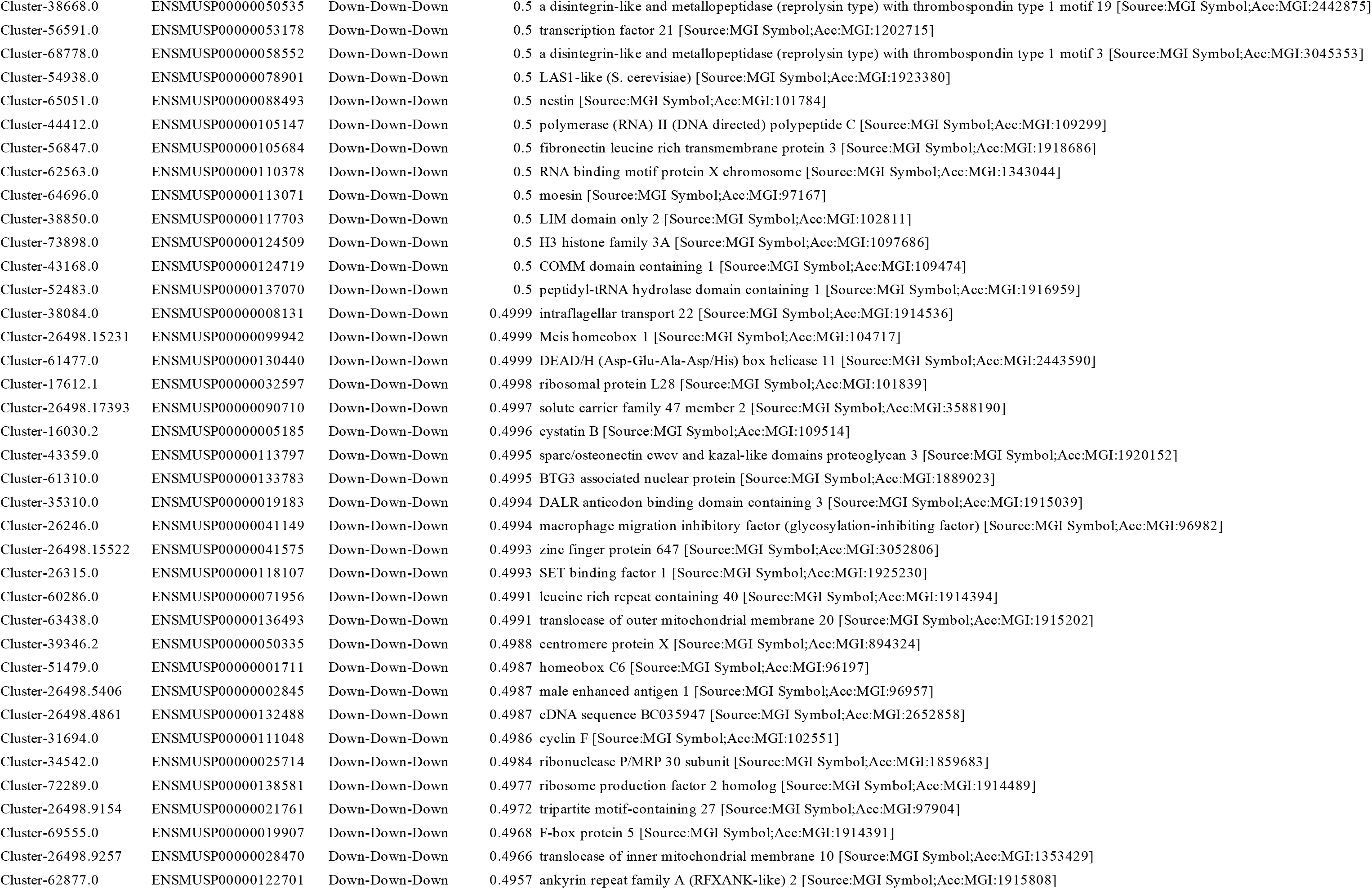
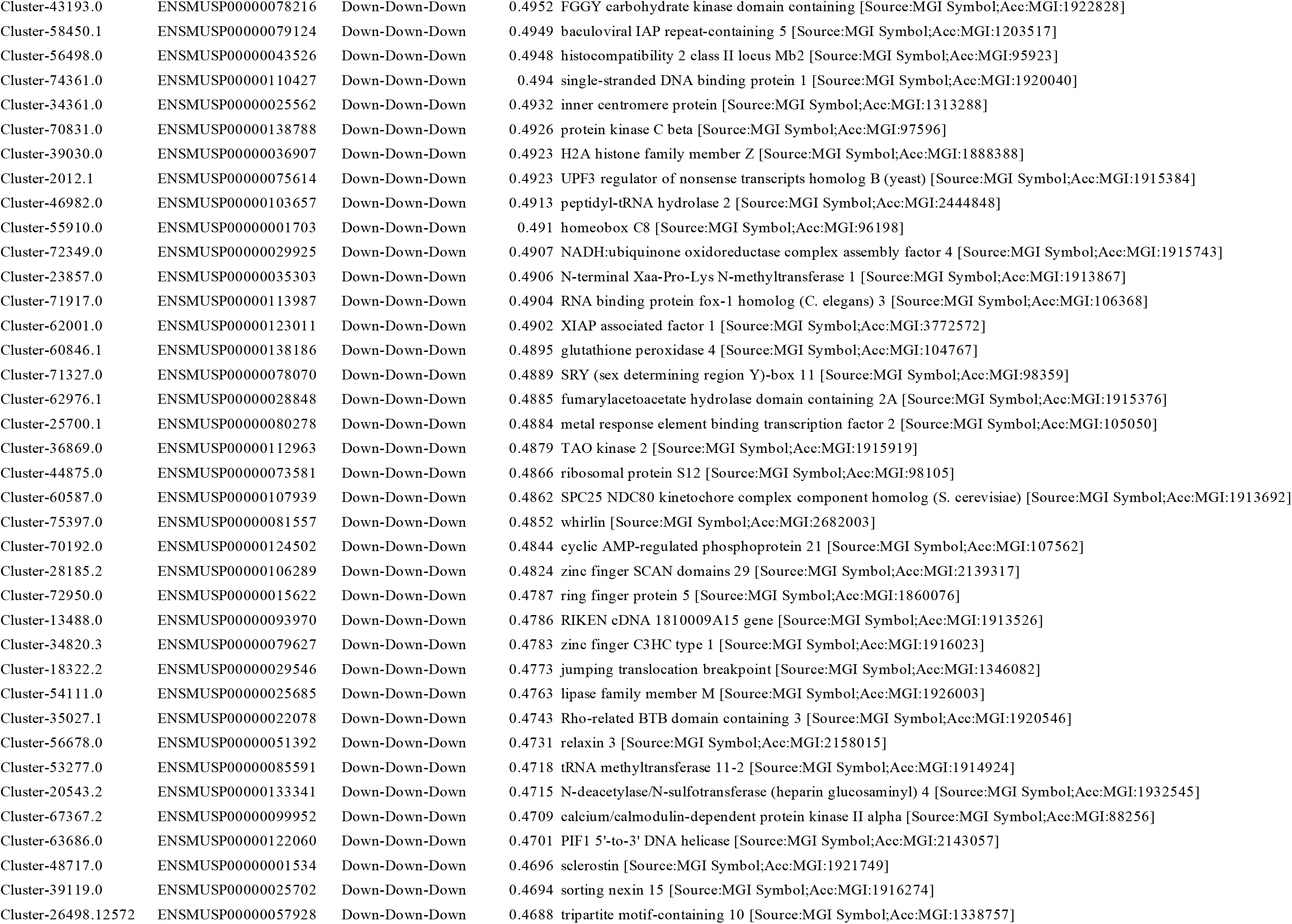
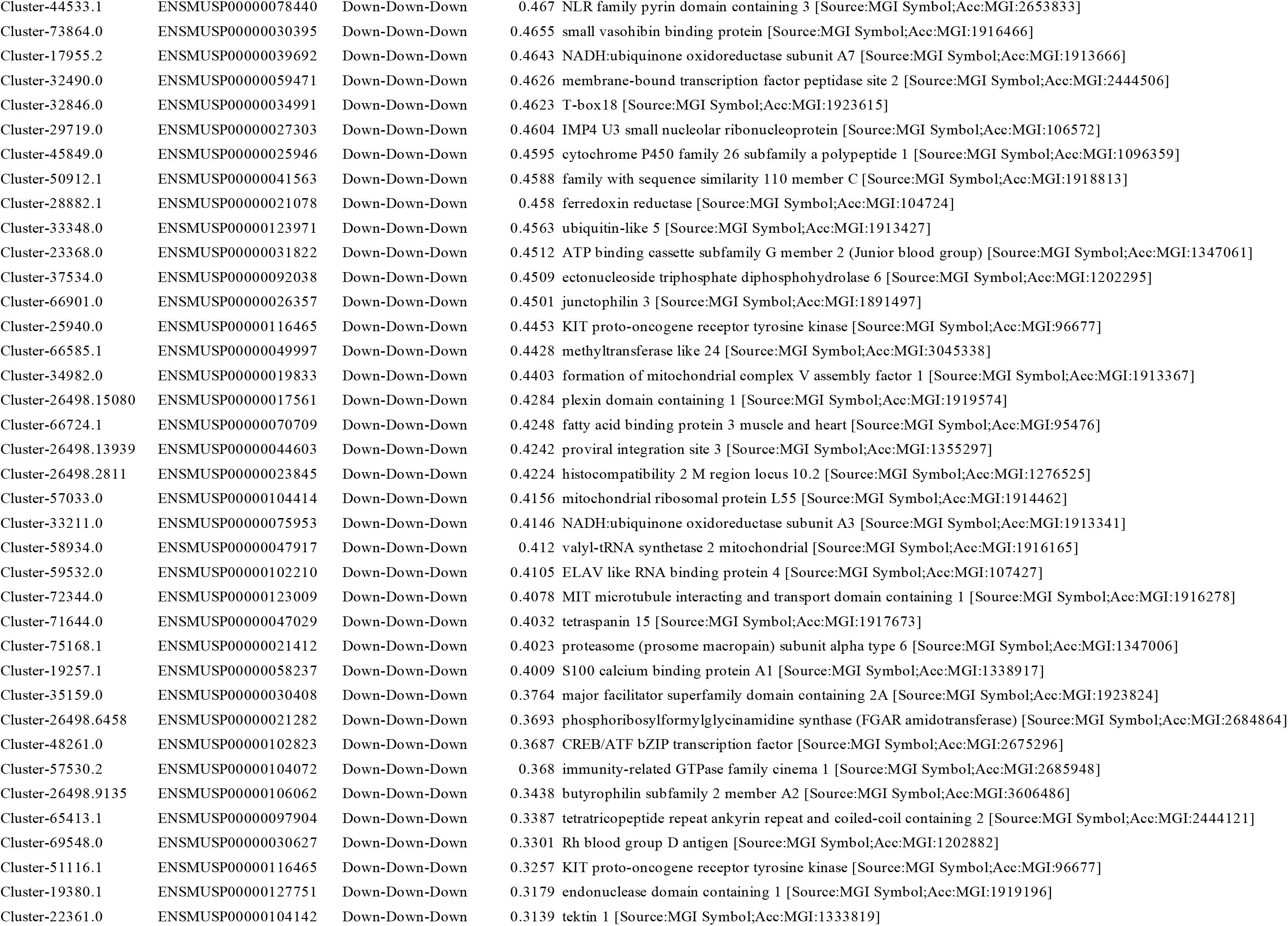
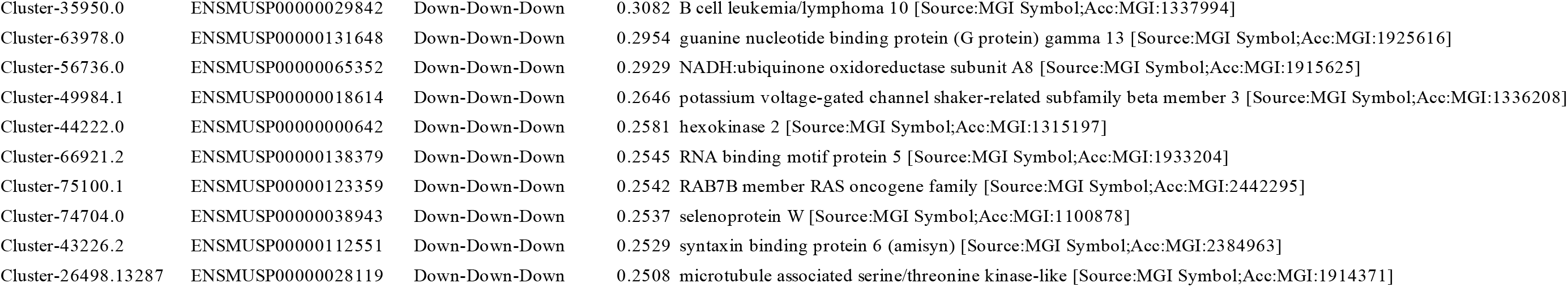
List of genes with categories of expresion pattern.

